# Identification of candidate susceptibility genes to *Puccinia graminis* f. sp. *tritici* in wheat

**DOI:** 10.1101/2021.01.23.427871

**Authors:** Eva C. Henningsen, Vahid Omidvar, Rafael Della Coletta, Jean-Michel Michno, Erin Gilbert, Feng Li, Marisa E. Miller, Chad L. Myers, Sean P. Gordon, John P. Vogel, Brian J. Steffenson, Shahryar F. Kianian, Cory D. Hirsch, Melania Figueroa

## Abstract

Wheat stem rust disease caused by *Puccinia graminis* f. sp. *tritici* (*Pgt*) is a global threat to wheat production. Fast evolving populations of *Pgt* limit the efficacy of plant genetic resistance and constrain disease management strategies. Understanding molecular mechanisms that lead to rust infection and disease susceptibility could deliver novel strategies to deploy crop resistance through genetic loss of disease susceptibility. We used comparative transcriptome-based and orthology-guided approaches to characterize gene expression changes associated with *Pgt* infection in susceptible and resistant *Triticum aestivum* genotypes as well as the non-host *Brachypodium distachyon*. We targeted our analysis to genes with differential expression in *T. aestivum* and genes suppressed or not affected in *B. distachyon* and report several processes potentially linked to susceptibility to *Pgt*, such as cell death suppression and impairment of photosynthesis. We complemented our approach with a gene co-expression network analysis to identify wheat targets to deliver resistance to *Pgt* through removal or modification of putative susceptibility genes.

## 1 Introduction

Stem rust caused by *Puccinia graminis* f. sp. *tritici* (*Pgt*) is one of the most devastating foliar diseases of wheat (*Triticum aestivum*) and barley (*Hordeum vulgare*). The economic relevance of this pathogen to food security is demonstrated by the impact of historical and recent epidemics (Singh et al., 2015; Pretorius et al., 2000; Olivera et al., 2015; Steffenson et al., 2017; Bhattacharya, 2017; Peterson, 2001). Consistent with its biotrophic lifestyle, *Pgt* develops an intricate relationship with its host in order to acquire nutrients and survive. Early stages of infection involve the germination of urediniospores (asexual spores) and host penetration through the formation of appressoria over stomata (Staples and Macko, 1984). As the fungus reaches the mesophyll cavity of the plant, it develops infection hyphae which penetrate plant cell walls and differentiate into specialized feeding structures, known as haustoria. Haustorial development takes place during the first 24 hours post-infection and is critical for colony establishment and sporulation that re-initiates the infection cycle (Harder and Chong, 1984). Similar to other plant pathogens, cereal rust infections involve the translocation of effectors to the plant cell as a mechanism to shut down basal defenses activated by PAMP triggered immunity (PTI) and manipulate host metabolism (Couto and Zipfel, 2016; Dodds and Rathjen, 2010). In rust fungi, the haustorium mediates the secretion of effectors, although the underlying molecular mechanism that facilitates this process is not known (Petre et al., 2014; Garnica et al., 2014). The plant targets of effectors and other plant genes that mediate compatibility and facilitate pathogen infection are often regarded as susceptibility (*S*) genes (van Schie and Takken, 2014; Lapin and Van den Ackerveken, 2013; Lo Presti et al., 2015).

To avoid infection by adapted pathogens, plants employ effector-triggered immunity (ETI) which is mediated by the recognition of effectors by nucleotide-binding domain leucine-rich repeat (NLR) receptors (Petre et al., 2014; Garnica et al., 2014; Dodds and Rathjen, 2010; Flor et al., 1971). These specific recognition events often induce localized cell death at infection sites (hypersensitive response, HR) which restrict pathogen growth. In wheat-rust interactions, ETI is manifested by the reduction or absence of fungal growth and sporulation (Periyannan et al, 2017). The use of NLR genes to provide crop protection was a critical component of the Green Revolution which diminished the impact of stem rust epidemics (Ellis et al., 2014). While this approach still contributes to the development of wheat cultivars with genetic resistance to stem rust, the durability of such resistant cultivars is hampered by the evolution of rust populations to avoid recognition by NLRs. Given the economic and environmental advantages of genetic disease control over chemical applications, the identification of alternative genetic sources of resistance are a priority for securing future wheat production. In this context, the discovery of *S* genes could have important translational applications for agriculture and potential durable disease control. Mutations in *S* genes, although often recessive, could shift a genotype to a non-suitable host due to alterations in initial recognition stages or loss of pathogen establishment requirements (van Schie and Takken, 2014; Lo Presti et al., 2015).

The genetic factors that contribute to wheat susceptibility to biotrophic pathogens such as rust fungi remain largely unknown. Numerous structural and physiological alterations have been observed in wheat-rust compatible interactions. At early infection stages, 4-6 days post-inoculation (dpi), the cytoplasm of infected mesophyll cells increases in volume and an extensive network of the endoplasmic reticulum is built near the haustorium (Bushnell, 1984). The nucleus of infected cells also increases in size and migrates towards the haustorium, and in some cases both structures appear in proximity. These observations suggest that plant cells undergo a massive transcriptional reprogramming to either accommodate rust colonization or initiate a cascade of plant defenses to prevent infection. In addition, many biotrophs are known to increase the ploidy of host cell nuclei near infection sites (Wildermuth, 2010). Advances in next generation sequencing and data mining bring new opportunities to deepen our understanding of plant-pathogen interactions and the relationship between plant metabolism and disease resistance or susceptibility. Several transcriptome profiling studies comparing compatible and incompatible wheat-rust interactions provide strong evidence for the complexity of these interactions (Bozkurt et al., 2010; Dobon et al., 2016; Chandra et al., 2016; Zhang et al., 2014; Yadav et al., 2016). Although *S* genes in rust pathosystems are largely unknown, several susceptibility factors to other plant pathogenic fungi have been identified in *Arabidopsis thaliana, Hordeum vulgare* (barley), and solanaceous plants (van Schie and Takken, 2014; Zaidi et al., 2018).

To expand our knowledge of wheat-rust interactions and identify candidate *S* genes to direct future functional studies, we conducted a comparative RNA-seq analysis of the molecular responses to *Pgt* in compatible and incompatible interactions. We included a susceptible genotype (W2691) of *Triticum aestivum* (bread wheat) and the same genotype containing the resistance gene *Sr9b*, which confers race-specific responses to various *Pgt* isolates (McIntosh et al., 1995). We also included the related grass species *Brachypodium distachyon*, which is recognized as a non-host to various cereal rust species (Kellogg, 2001; Figueroa et al., 2013, 2015; Ayliffe et al., 2013; Omidvar et al., 2018; Gilbert et al., 2018; Bettgenhaeuser et al., 2018, 2014). As part of our analysis, we examined the expression profiles of *T. aestivum* and *B. distachyon* orthologs of several known *S* genes in *Arabidopsis thaliana, H. vulgare*, as well as other characterized *S* genes in *T. aestivum*, and identified groups of genes co-regulated with these *S* gene candidates. In conclusion, this study provides an overview of global expression changes associated with failure or progression of *Pgt* infection in *T. aestivum* and *B. distachyon* and insights into the molecular processes that define disease incompatibility.

## 2 Materials and Methods

### 2.1 Plant and fungal materials

Two near-isogenic lines of *T. aestivum*, W2691 (Luig and Watson, 1972) and W2691 carrying the *Sr9b* gene (referred to onward as W2691*+Sr9b*, U.S. National Plant Germplasm System Accession Identifier: CI 17386) and the *B. distachyon* Bd21-3 inbred line (Vogel and Hill, 2008) were used in this study. *T. aestivum* and *B. distachyon* seeds were received from the USDA-ARS Cereal Disease Laboratory (CDL) St. Paul, MN, USA and the USDA-ARS Plant Science Unit, St. Paul, MN, USA, respectively. The fungal isolate *Puccinia graminis* f. sp. *tritici (Pgt)* (isolate # CDL 75-36-700-3 race SCCL) (Duplessis et al., 2011) was obtained from the USDA-ARS CDL.

### 2.2 *Pgt* infection of *T. aestivum* and *B. distachyon* genotypes

*B. distachyon* seeds were placed in petri dishes with wet grade 413 filter paper (VWR International) at 4°C for five days and germinated at room temperature for three days before sowing to synchronize growth with wheat plants which did not require stratification. Seeds of both wheat and *B. distachyon* were sown in Fafard® Germination Mix soil (Sun Gro Horticulture, Agawam, MA, U.S.A.). All plants were grown in growth chambers with a 18/6 hour light/dark cycle at 21/18°C light/dark and 50% relative humidity. Urediniospores of *Pgt* were activated by heat-shock treatment at 45°C for 15 minutes and suspended in Isopar M oil (ExxonMobil) at 10 mg/mL concentration. Inoculation treatments consisted of 50 µl of spore suspension per plant, whereas mock treatments consisted of 50 µl of oil per plant. Fungal and mock inoculations were conducted on seven-day old wheat plants (first-leaf stage) and twelve-day old *B. distachyon* plants (three-leaf stage). After inoculations, plants were kept for 12 hours in mist chambers with repeated misting for 2 minutes every 30 minutes and returned to growth chambers under the previously described conditions.

### 2.3 Analysis of fungal colonization and growth

At 2, 4, and 6 dpi *T. aestivum* and *B. distachyon* leaves were sampled and cut into 1 cm sections before staining with Wheat Germ Agglutinin Alex Fluor® 488 conjugate (WGA-FITC; ThermoFisher Scientific) following previously described procedures (Omidvar et al., 2018). Time points to represent stages of *Pgt* infection were selected based on previous characterization (Figueroa et al., 2013; Figueroa et al., 2015). To determine the level of fungal colonization, the percentage of urediniospores that germinated (GS), formed an appressorium (AP), established a colony (C), and differentiated a sporulating colony (SC) were visualized using a fluorescence microscope (Leica model DMLB; 450-490 nM excitation). The progression of fungal growth was recorded for 100 infection sites for each of the three biological replicates. Genomic DNA was extracted from *T. aestivum* (three infected primary leaves) and *B. distachyon* (three infected secondary leaves) using the DNeasy Plant Mini Kit (Qiagen) and were standardized to a 10 ng/µl concentration. The ITS regions were amplified by qPCR using ITS-specific primers provided by the Femto(tm) Fungal DNA Quantification Kit (Zymo Research) to quantify the relative abundance of fungal DNA following the manufacturer’s recommendations for the three biological replicates. The *GAPDH* housekeeping gene from each species was used as an internal control to normalize fungal DNA quantities (Omidvar et al., 2018).

### 2.4 RNA isolation, purification, and sequencing

Infected and mock treated primary leaves from W2691 and W2691+*Sr9b* and secondary leaves from Bd21-3 were collected at 2, 4, and 6 dpi. For each of the three biological replicates, three infected leaves were pooled for RNA extraction using the RNeasy Plant Mini Kit (Qiagen). Subsequently, stranded-RNA libraries were constructed, and 125 bp paired-end reads were sequenced on an Illumina HiSeq(tm) 2500 instrument at the University of Minnesota Genomics Center. On average, more than 10 million reads were generated per time point in each of the previously listed plant-rust interactions (**Table S1**).

### 2.5 Alignment of reads to the *T. aestivum* and *B. distachyon* reference genomes

Short reads and low-quality bases were trimmed using cutadapt v1.18 (Martin, 2011) with the following parameters: minimum-length 40, quality-cutoff 30, and quality-base=33. Subsequently, W2691 and W2691+*Sr9b* reads were mapped to the *T. aestivum* cv. Chinese Spring reference genome IWGSC RefSeq v1.0 (Alaux et al., 2018) and Bd21-3 reads were mapped to the Bd21-3 reference genome from the Joint Genome Institute (*B. distachyon* Bd21-3 v1.1 DOE-JGI, http://phytozome.jgi.doe.gov/). Read mapping was conducted using STAR v2.5.3 (Dobin et al., 2012) set for two-pass mapping mode with the following parameters: twopassMode Basic and outSAMmapqUnique 20.

### 2.6 Expression profiling and identification of differentially expressed genes

Reads were mapped to *T. aestivum* and *B. distachyon* gene features using htseq v.0.11.0 to obtain count values (Anders et al., 2015). Normalized read counts and differential expression (DE) analysis were performed with DESeq2 v1.28.1 (Love et al., 2014). Genes with a |log2 fold change| ≥ 1.5 and a *p*-value < 0.05 were identified as differentially expressed genes (DEGs).

### 2.7 Gene ontology analysis

Gene ontology (GO) terms were obtained from GOMAP track data for *T. aestivum* (Alaux et al., 2018) and previously published data for *B. distachyon* (*Brachypodium distachyon* Bd21-3 v1.2 DOE-JGI, http://phytozome.jgi.doe.gov/) annotation files. GO terms in wheat and *B distachyon* were mapped to the GOslim plant subset using OWLTools with the command owltools ╌map2slim (https://github.com/owlcollab/owltools). GO enrichment analysis for DEGs was performed using the topGO R package using the “weight01” algorithm and fisher test statistic (Alexa and Rahnenfuhrer, 2020). Enriched terms were considered significant with a Fisher test *p*-value < 0.01 (**Table S2**). Enrichment analyses using the GOslim subset were performed on all differentially expressed wheat and *B. distachyon* genes, as well as on genes within the *S*-gene orthologs clusters. Enrichment analysis with the full GO set was only performed on the differentially expressed *T. aestivum* and *B. distachyon* genes using the same methods described above.

### 2.8 Orthology analysis

Protein sequences from *S* genes of interest (**Table S3**) as cited in original publications as reviewed by van Schie and Takken (2014) were cross-checked using gene name and synonym information and the Basic Local Alignment Search Tool (BLAST) functions in the TAIR gene search database (https://www.arabidopsis.org/index.jsp), EnsemblPlants (https://plants.ensembl.org/index.html), UniPro (https://www.uniprot.org/), and the IPK blast server (https://webblast.ipk-gatersleben.de/barley_ibsc/). OrthoFinder version 2.4.0 (Emms and Kelly, 2019) was used to identify orthologs between *A. thaliana* (https://phytozome.jgi.doe.gov/pz/portal.html#!info?alias=Org_Athaliana), *H. vulgare* (http://floresta.eead.csic.es/rsat/data/genomes/Hordeum_vulgare.IBSCv2.36/genome/Hordeum_vulgare.IBSCv2.36.pep.all.fa), *T. aestivum* (annotation version 1.1 https://urgi.versailles.inra.fr/download/iwgsc/IWGSC_RefSeq_Annotations/v1.1/iwgsc_refseqv1.1_genes_2017July06.zip), and *B. distachyon* (annotation version 1.2 https://phytozome-next.jgi.doe.gov/info/BdistachyonBd21_3_v1_2) proteins. For genes in *A. thaliana, H. vulgare*, and *T. aestivum* with multiple isoforms, perl scripts for each species were used to retain only the longest representative transcript for use in the orthology analysis (https://github.com/henni164/stem_rust_susceptibility/longest_transcript/perl). The longest transcript file for *B. distachyon* (BdistachyonBd21_3_537_v1.2.protein_primaryTranscriptOnly.fa) was obtained from Phytozome. The default settings of OrthoFinder were used, and orthologs of the four species were obtained in a single run. The URGI BLAST tool (https://wheat-urgi.versailles.inra.fr/Seq-Repository/BLAST) was used to identify candidates for missing subgenome representatives.

### 2.9 Protein sequence phylogenetic analysis

Using the longest protein sequence from known *S* genes in *A. thaliana* (Lamesch et al., 2011) and *H. vulgare* (Howe et al., 2019), as well as the longest protein sequences from the orthologous candidate *S* genes in *T. aestivum* and *B. distachyon* (**Table S3**), phylogenetic trees were constructed to examine the relationship of ortholog families using the web-based tool NGPhylogeny (Lemoine et al., 2019). Default parameters for the FastME one-click workflow were used for MAFFT alignment, BMGE curation, and FASTME tree inference (https://ngphylogeny.fr/documentation). A R script using the packages ggplot2, ggtree, and ape was used to generate visualizations of the generated phylogenetic trees (Wickham, 2016; Yu et al., 2017; Paradis and Schliep, 2018).

### 2.10 Gene co-expression network analysis

Individual gene co-expression networks (GCNs) were constructed and analyzed for *T. aestivum* W2691, W2691+*Sr9b* and *B. distachyon* Bd21-3 genotypes using the python package Camoco (Schaefer et al., 2018). To build each network, all three independent RNA-seq replicates from all three time points (2, 4, and 6 dpi) of infected and mock-inoculated treatments were used. HTSeq read counts were converted to FPKM values for Camoco compatibility, and subjected to inverse hyperbolic sine transformation normalized against median FPKMs across all samples. Genes with coefficient of variation < 0.1 across all samples or without a single sample having an expression above 0.5 FPKM were removed from analysis. Additionally, genes with a FPKM value > 0.001 across 60% of samples were included in network analyses. Pearson correlation metrics between all gene pairs were calculated and subjected to Fisher transformation to generate Z-scores with a cutoff of Z >= 3 to allow comparisons between networks (Huttenhower et al., 2006). Finally, correlation metrics were used to build weighted gene co-expression networks. Clusters containing susceptibility gene orthologs were visualized using ggplot2 (Wickham, 2016), ggnetwork (Briatte, 2020), sna (Butts, 2019), and network (Butts, 2015) R packages.

### 2.11 Data availability

Sequence data was deposited in NCBI under BioProject PRJNA483957 (**Table S1**). Unless specified otherwise, supplemental tables, scripts and files for analysis and visualizations are available at https://github.com/henni164/stem_rust_susceptibility.

## 3 Results

### 3.1 *T. aestivum* and *B. distachyon* differ in susceptibility to *Pgt*

We compared the infection and colonization of *Pgt* in two *T. aestivum* isogenic lines that were susceptible (W2691) or resistant (W2691+*Sr9b*) to *Pgt* as well as in the non-host *B. distachyon* Bd21-3. Symptom development upon infection was consistent with previous observations reporting susceptibility of W2691 and W2691+*Sr9b* mediated resistance (intermediate) to race SCCL (**Figure 1A, B)** (Zambino et al., 2000). Susceptibility was manifested by formation of large sporulating pustules in W2691, while small pustules surrounded by a chlorotic halo were characteristic of *Sr9b* mediated-resistance at 6 days post-inoculation (dpi). Susceptibility differences between W2691 and W2691+*Sr9b* were evident at 6 dpi as formation of fungal colonies was present in both genotypes, but colony sizes were larger in W2691 than W2691+*Sr9b* (**Figure 1**). *B. distachyon* supports the formation of colonies that are smaller than those in the resistant *T. aestivum* line W2691+*Sr9b* with no visible macroscopic symptoms observed at 6 dpi (**Figure 1C**). To monitor the progression of fungal growth, we quantified the percentage of germinated urediniospores (GS), and interaction sites displaying the formation of appressoria (AP), colony formation (C), and colony sporulation (SC) at 2, 4, and 6 dpi using microscopy (**Figure 1D**). The germination frequency (∼95%) was similar between all three genotypes tested (ANOVA test, *p* > 0.05). The percentage of interaction sites showing appressorium formation (AP) was higher in wheat than in *B. distachyon* at 4 dpi (ANOVA test, *p* ≤ 0.035). The genotype W2691 displayed the highest percentage of interaction sites showing colony formation at 4 and 6 dpi (ANOVA test, *p* ≤ 0.002), and sporulation at 6 dpi (ANOVA test, *p* ≤ 0.0015). In contrast, a smaller number of rust colonies formed in *B. distachyon*, and these colonies did not show signs of sporulation. To estimate rust colonization levels on *T. aestivum* and *B. distachyon*, we quantified the abundance of fungal DNA in infected leaves at 2, 4, and 6 dpi (**Figure 1E**). Rust colonization among all genotypes was not significantly different at 2 dpi (ANOVA test, *p* > 0.05); however, there was a trend at 4 and 6 dpi for higher rust colonization in W2691 than in W2691+*Sr9b* and *B. distachyon* (ANOVA test, *p* > 0.05).

**Figure 1.**
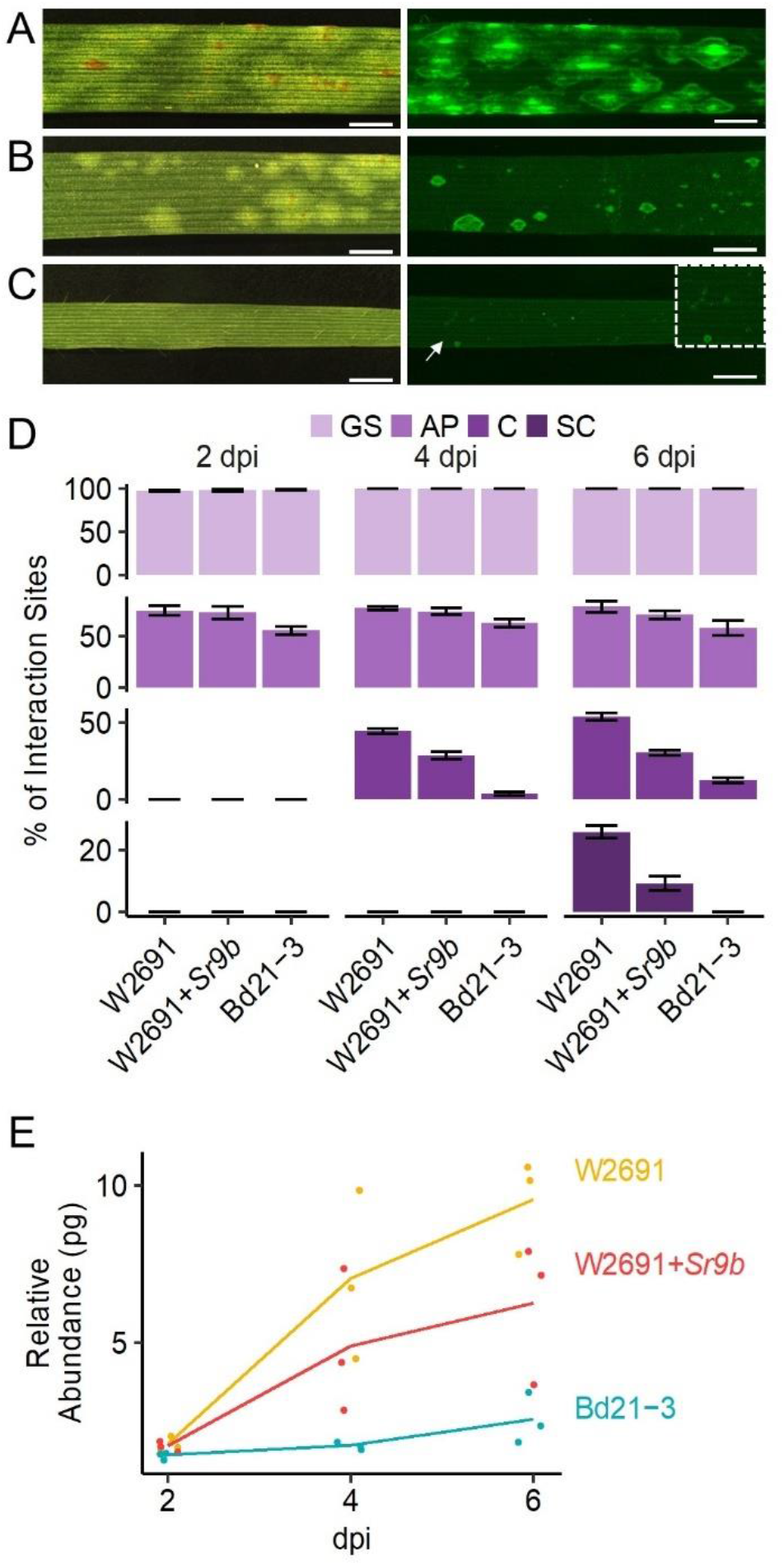
Infection of *T. aestivum* and *B. distachyon* genotypes with *P. graminis* f. sp. *tritici* race SCCL. (**A-C**) Development of disease symptoms (left) and fungal colonization (right) at 6 dpi. (**A**) W2691 (susceptible wheat line). (**B**) W2691+*Sr9b* (intermediate resistant wheat line). (**C**) *B. distachyon* Bd21-3 line (non-host). The white arrow and the white box indicate the area which was enlarged for better visualization of colonies. Scale bars indicate 2 mm. (**D**) Percentage of fungal infection sites which showed germinated urediniospores (GS), appressorium formation (AP), colony establishment (C), and sporulating colony (SC). Error bars represent the standard error of three independent biological replicates. (**E**) Fungal DNA abundance in infected W2691, W2691+*Sr9b*, and Bd21-3 genotypes as measured using qPCR. The points show the sample values and the lines represents the mean of the samples.

### 3.2 Putative biological processes associated with *in planta* responses to *Pgt*

The transcriptome profiles of *T. aestivum* (W2691 and W2691+*Sr9b*) and *B. distachyon* (Bd21-3) in response to *Pgt* infection at 2, 4, and 6 dpi were examined using RNA-seq expression profiling (**Table S1**). Differential expression analysis was used to compare responses to rust infection relative to the baseline mock treatments. Overall, the number of differentially expressed genes (DEGs) increased in W2691, W2691+*Sr9b*, and Bd21-3 over the course of infection (**Table 1**). Between 11-12.9% of *T. aestivum* genes were differentially expressed at 6 dpi, whereas in Bd21-3 only 6.2% were differentially expressed. We conducted a GOslim enrichment analysis on up- and down-regulated DEGs for each interaction at the infection time points **(Figure 2**). At 2 dpi, W2691 and W2691+*Sr9b* had only a few GOslim terms enriched in either up- or down-regulated DEGs. At 4 dpi, greater similarities between the *T. aestivum* genotypes emerged with very similar enrichment patterns in GOslim terms. The similarity of GOslim term enrichment continued at 6 dpi, with W2691 and W2691+*Sr9b* having nearly identical enrichment patterns. W2691+*Sr9b* had one additional term enriched in both up-regulated (cytoplasm, GO:0005737) and down-regulated (chromatin binding, GO:0003682) genes. Compared to the two *T. aestivum* genotypes, Bd21-3 had fewer terms enriched across all three timepoints and only a few terms were in common with W2691 and W2691+*Sr9b* (i.e., extracellular region (GO:0005576), DNA-binding transcription factor activity (GO:0003700). Bd21-3 had several unique terms in both up- and down-regulated categories, among them mitochondrion (GO:0005739), transporter activity (GO:0005215), catalytic activity (GO:0003824), and DNA binding (GO:0003677) were upregulated, while intracellular (GO:0005622), DNA-binding transcription factor activity (GO:0003700), catalytic activity (GO:0003824), and DNA binding (GO:0003677) were downregulated. The full GO set also demonstrated clear differences between the *T. aestivum* genotypes and Bd21-3. Photosynthesis-related terms such as chloroplast photosystem I and II (GO:0030093 and GO:0030095), photosystem II antenna complex (GO:0009783), and PSII associated light-harvesting complex II (GO:0009517) were overrepresented at 4 and 6 dpi in W2691 and W2691+*Sr9b*, but not in Bd21-3 (**Table S2**). In addition, Bd21-3 only had enrichment in 11 terms across the cellular component (CC), biological process (BP), and molecular function (MF) categories compared to the terms enriched 741 across the three categories in W2691 and W2691+*Sr9b* (**Table S2**). Overall, this analysis highlights how the molecular and genetic responses of Bd21-3 to *Pgt* differ from those in W2691 and W2691+*Sr9b* over the course of the experiment.

**Table 1.** Differentially expressed genes in *T. aestivum* and *B. distachyon* in response to *P. graminis* f. sp. *tritici* infection.

|  | **2 dpi** |  | **4 dpi** |  | **6 dpi** |  |
| --- | --- | --- | --- | --- | --- | --- |
| **Genotype** | **up** | **down** | **up** | **down** | **up** | **down** |
| W2691 | 278 | 577 | 2,887 | 2,614 | 6,659 | 7,241 |
| W2691+*Sr9b* | 747 | 110 | 3,835 | 1,832 | 6,397 | 5,471 |
| Bd21-3 | 200 | 437 | 559 | 1,419 | 739 | 1,665 |

\* Total number of wheat genes: 107,891; total number of *B. distachyon* genes: 39,068. Within- genotype comparisons used mock treatments as the baseline.

**Figure 2.**
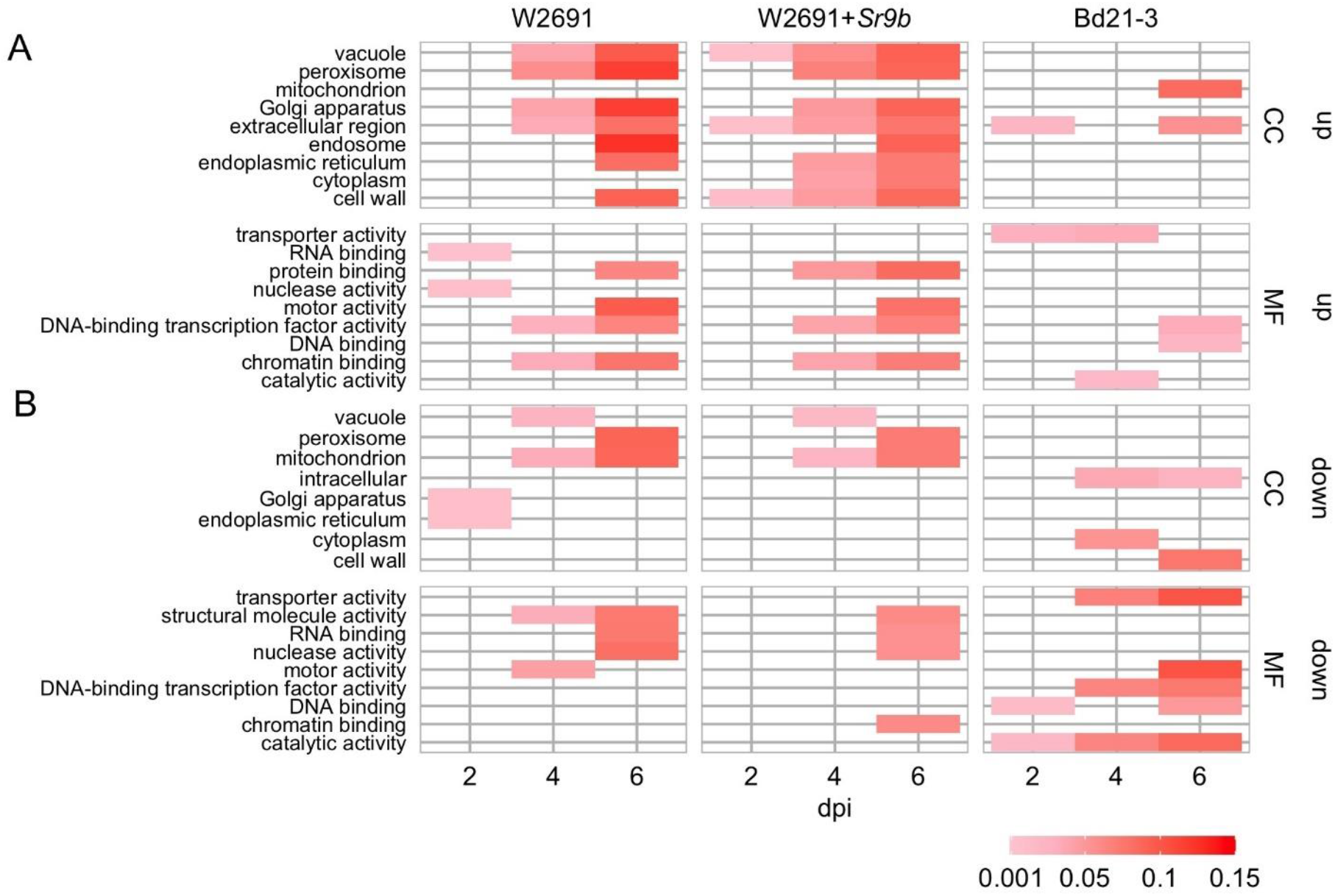
GOslim enrichment analysis of differentially expressed (DE) genes in mock vs inoculated *T. aestivum* (W2691 and W2691+*Sr9b*) and *B. distachyon* (Bd21-3) genotypes across three time points (bottom x-axis) upon infection with *P. graminis* f. sp. *tritici*. (**A**) Enrichment of plant GOslim terms of upregulated (up) DE genes and (**B**) downregulated (down) DE genes. The y-axis shows plant GO slim terms separated by category: cellular component (CC) and molecular function (MF). The scale represents the proportion of genes annotated with each GO term to all the genes tested.

### 3.3 Differential regulation of candidate orthologous susceptibility (S) genes in *T. aestivum* and *B. distachyon* upon *Pgt* infection

Various *S* genes have been previously characterized or postulated in several species, including *A. thaliana* and *H. vulgare* (Büschges et al., 1997; Chen et al., 2007; Chen et al., 2010; Low et al., 2020), and this knowledge has allowed us to further understand molecular plant-microbe interactions. With an interest in identifying potential *S* genes in *T. aestivum* as well as creating resources to enable future studies, we designed an experimental workflow based on the identification of known *S* gene orthologs, gene expression comparisons and co-expression network analysis (**Figure 3**). A curated set of previously characterized or postulated *S* genes as summarized by Schie and Takken (2014) was narrowed down by selecting genes in *A. thaliana*, and *H. vulgare*, and eliminating *S* genes that were discovered or characterized for viruses or necrotrophic fungi, leaving 112 potential candidate *S* genes to examine (**Table S3**). We then conducted an orthology analysis using all *H. vulgare, A. thaliana, B. distachyon*, and *T. aestivum* transcripts to identify orthogroups of longest transcript of all genes. Orthogroups were constructed from 211,973 genes across these species (**Table S3**). A total of 182,206 genes were assigned to 29,420 orthogroups, the largest of which (OG0000000) contained 211 genes. Of the total genes, 92,913 (86%) wheat, 31,334 *B. distachyon* (80%), 34,075 barley (91%), and 23,883 *A. thaliana* (87%) genes were assigned to orthogroups. We identified 91 of the reported *S* genes from *A. thaliana* and *H. vulgare* across 70 orthogroups, that also consisted of at least one *T. aestivum* gene and one *B. distachyon* gene (**Table S4**). These genes from *T. aestivum* and *B. distachyon* were selected as *S* gene orthologs. A total of 29,767 genes (orthogroup OG0029421 to OG0059187) were assigned groups with only one member (singleton orthogroups) (**Table S5**).

**Figure 3.**
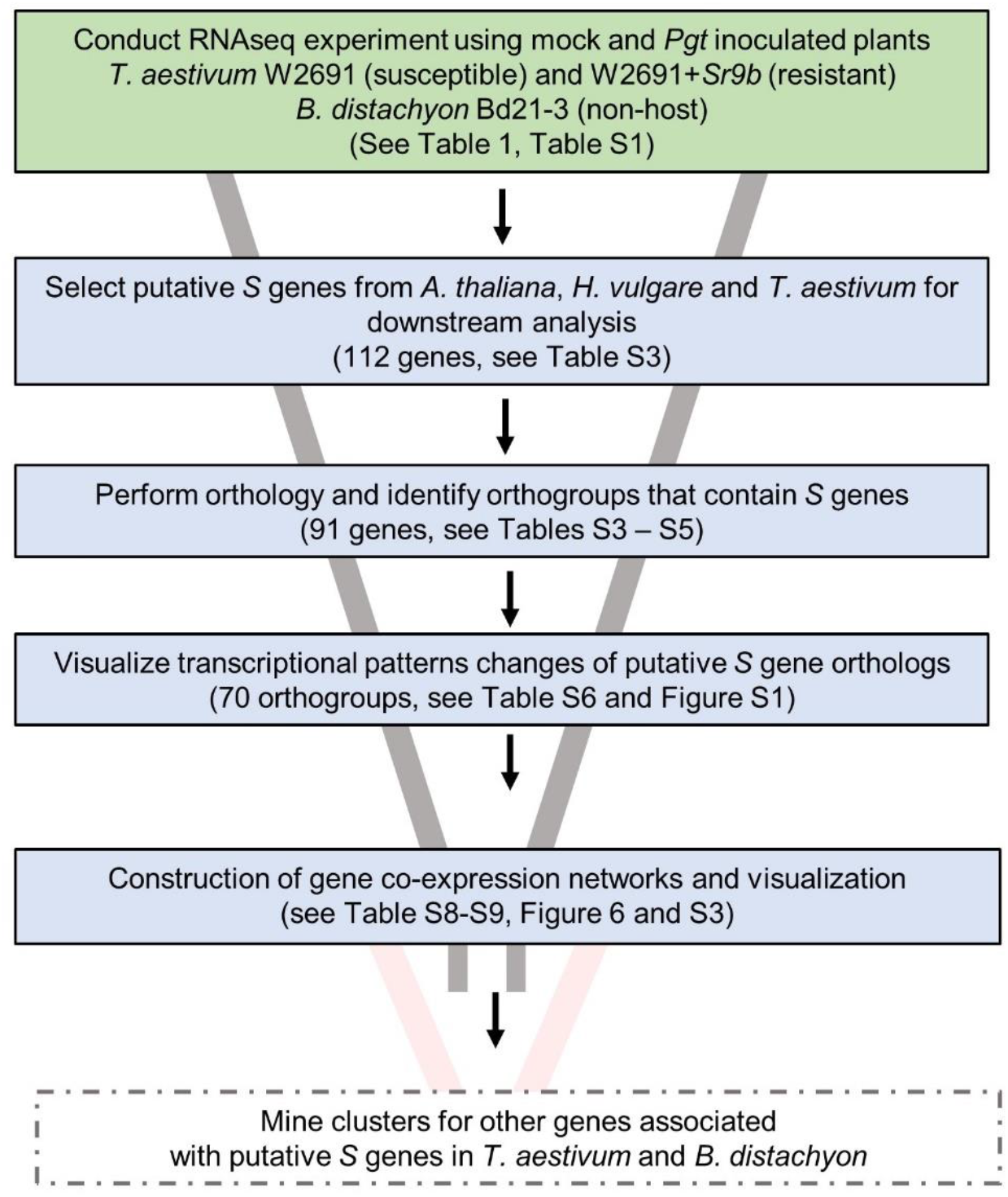
Experimental workflow used to identify candidates of *S* genes that contribute to infection of *T. aestivum* by *P. graminis* f. sp. *tritici*. Solid box outlines indicate work completed in this publication. Future work is indicated by dashed box outlines.

The gene expression patterns of *S* gene orthologs in *T. aestivum* and *B. distachyon* were used to identify which orthologs may act as susceptibility factors (**Figure 3, Table S6**). The selection criterion was applied to include DEGs that showed a progressive increase in log2 fold change (mock vs infected, |log2 fold change| ≥ 1.5 and a *p*-value < 0.05) in W2691 or in both W2691 and W2691+*Sr9b*, but the corresponding orthologs in *B. distachyon* and/or W2691+*Sr9b* showed a decrease or no change, as observed in various systems (Chen et al., 2010; Pessina et al., 2014). The assumption is that *S* genes will be up-regulated during infection when the pathogen reaches the sporulation stage (e.g., in a susceptible or intermediate resistant host represented by W2691 and W2691+*Sr9b*, respectively) but with a low or no regulatory change in a non-host (Bd21-3). Expression data for all genes can be found in **Table S6** in association with orthogroup number. Most genes in the 70 orthogroups did not demonstrate major changes in expression over the course of the experiment (**Figure S1**), including the orthogroup OG0001703, which contains the *Mlo* (*Mildew locus O*) alleles and orthologous sequences. Eight orthogroups that demonstrated these expression patterns were chosen for further analysis; these included ortholog genes for *AGD2* (*aberrant growth and death 2*), *BI-1* (*BAX inhibitor-1*), *DMR6* (*downy mildew resistance 6*), *DND1* (*defense, no death*), *FAH1* (*fatty acid hydroxylase 1*), *IBR3* (*IBA response 3*), *VAD1* (*vascular associated death 1*), and *WRKY25* (*WRKY DNA binding protein 25*) (**Figure 4, Table 2, Table S7**). Among the eight susceptibility orthogroups, *T. aestivum* orthologs of *BI-1, DMR6*, and *WRKY25* showed the greatest increase in fold change (**Table S7**) in either W2691 or W2691+*Sr9b*, particularly at 6 dpi (**Figure 4**). The gene ortholog of *DND1* displayed a higher fold change in W2691 than in W2691+*Sr9b*.

**Table 2.**
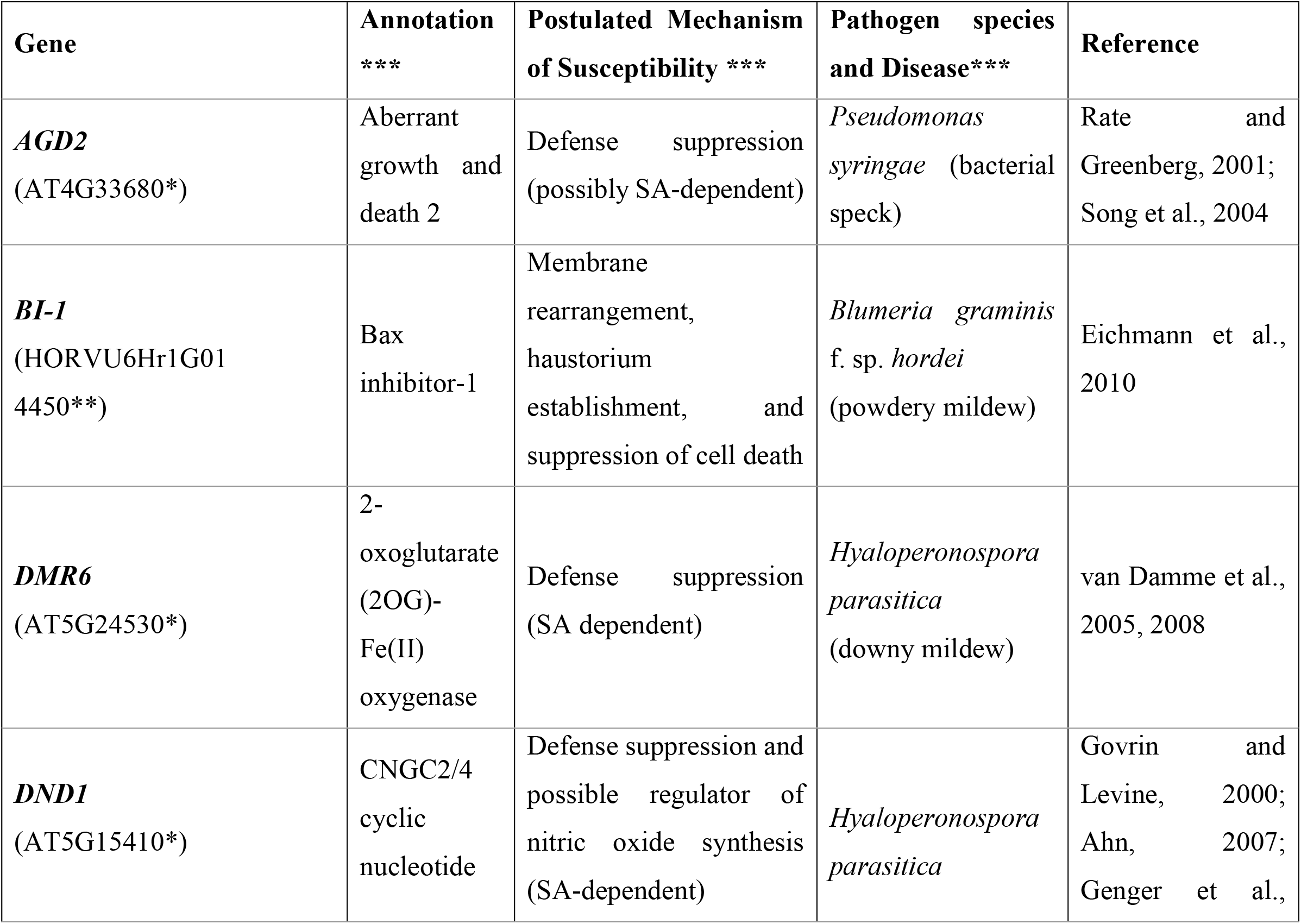
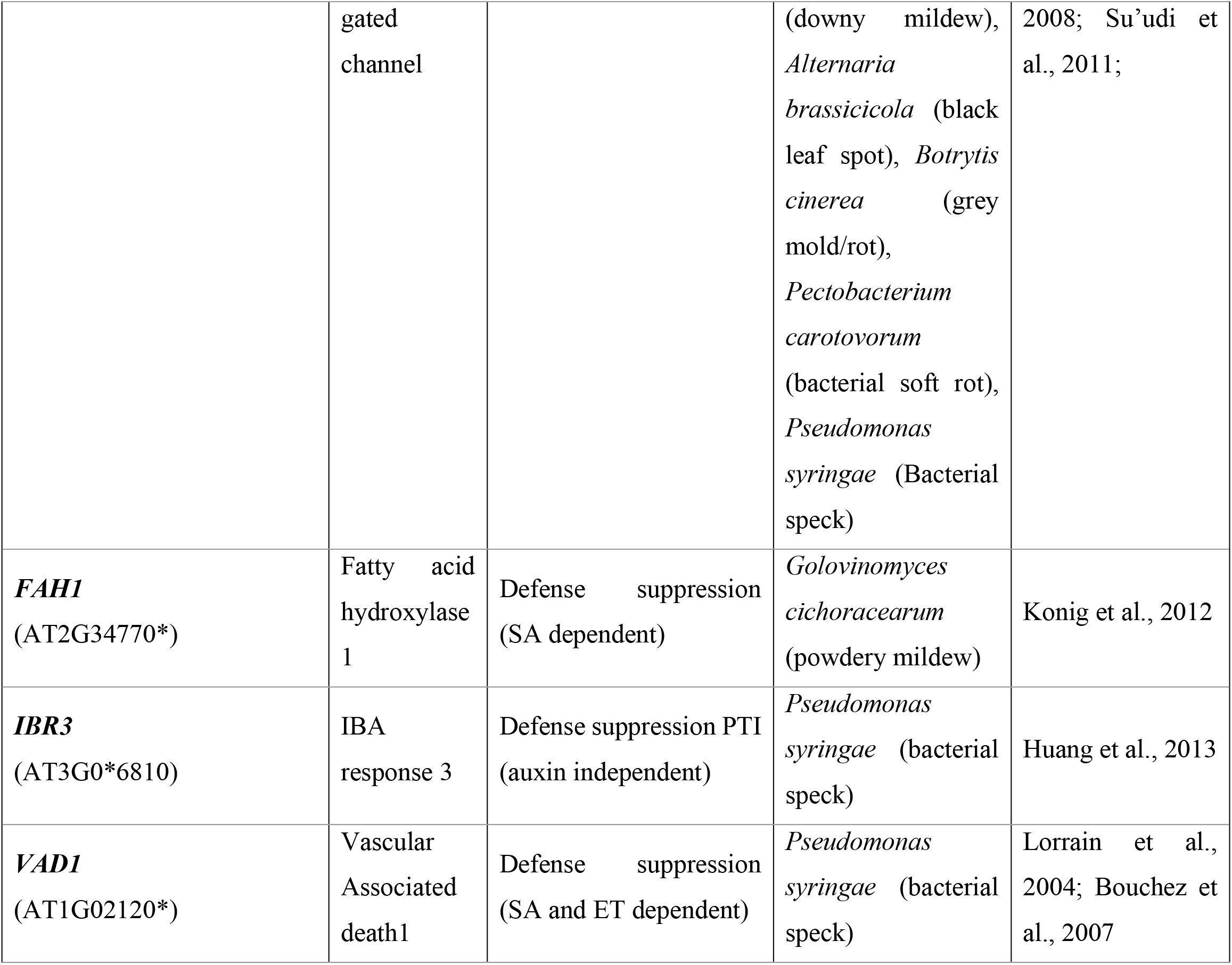
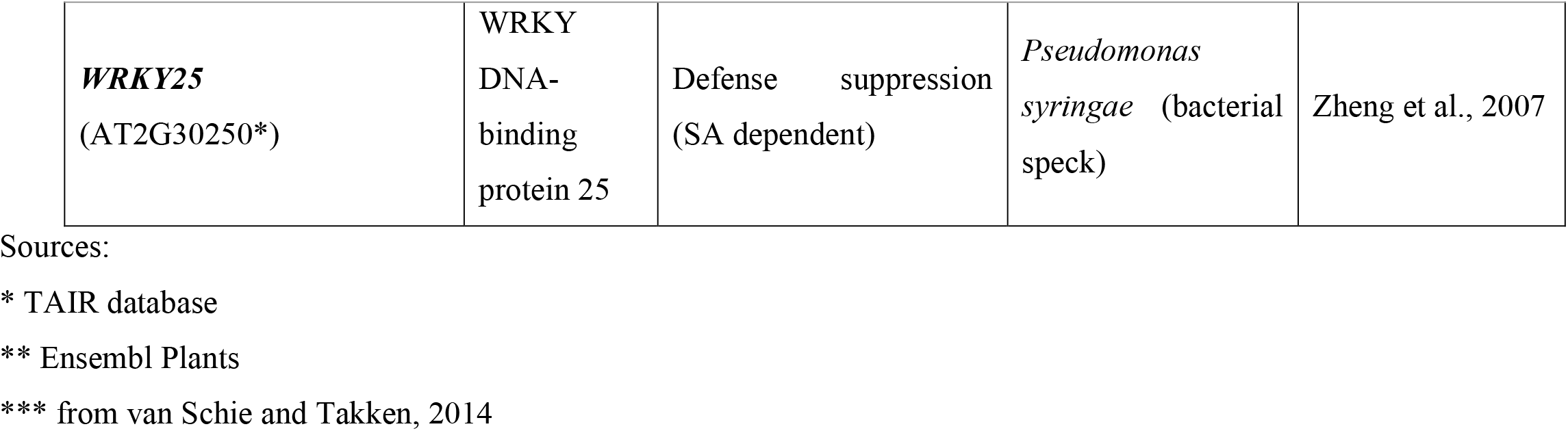
List of *S* genes explored through the gene expression analysis.

| Gene | Annotation<br>*** | Postulated Mechanism<br>of Susceptibility *** | Pathogen species<br>and Disease*** | Reference |
| --- | --- | --- | --- | --- |
| ***AGD2***<br>(AT4G33680*) | Aberrant<br>growth and<br>death 2 | Defense suppression<br>(possibly SA-dependent) | *Pseudomonas*<br>*syringae* (bacterial<br>speck) | Rate and<br>Greenberg, 2001;<br>Song et al., 2004 |
| ***BI-1***<br>(HORVU6Hr1G01<br>4450**) | Bax<br>inhibitor-1 | Membrane<br>rearrangement,<br>haustorium<br>establishment, and<br>suppression of cell death | *Blumeria graminis*<br>f. sp. *hordei*<br>(powdery mildew) | Eichmann et al.,<br>2010 |
| ***DMR6***<br>(AT5G24530*) | 2-oxoglutarate<br>(2OG)-<br>Fe(II)<br>oxygenase | Defense suppression<br>(SA dependent) | *Hyaloperonospora*<br>*parasitica*<br>(downy mildew) | van Damme et al.,<br>2005, 2008 |
| ***DND1***<br>(AT5G15410*) | CNGC2/4<br>cyclic<br>nucleotide | Defense suppression and<br>possible regulator of<br>nitric oxide synthesis<br>(SA-dependent) | *Hyaloperonospora*<br>*parasitica* | Govrin and<br>Levine, 2000;<br>Ahn, 2007;<br>Genger et al., |
|  | gated<br>channel |  | (downy mildew),<br>*Alternaria*<br>*brassicicola* (black<br>leaf spot), *Botrytis*<br>*cinerea* (grey<br>mold/rot),<br>*Pectobacterium*<br>*carotovorum*<br>(bacterial soft rot),<br>*Pseudomonas*<br>*syringae* (Bacterial<br>speck) | 2008; Su'udi et<br>al., 2011; |
| ***FAH1***<br>(AT2G34770*) | Fatty acid<br>hydroxylase<br>1 | Defense suppression<br>(SA dependent) | *Golovinomyces*<br>*cichoracearum*<br>(powdery mildew) | Konig et al., 2012 |
| ***IBR3***<br>(AT3G0*6810) | IBA<br>response 3 | Defense suppression PTI<br>(auxin independent) | *Pseudomonas*<br>*syringae* (bacterial<br>speck) | Huang et al., 2013 |
| ***VAD1***<br>(AT1G02120*) | Vascular<br>Associated<br>death1 | Defense suppression<br>(SA and ET dependent) | *Pseudomonas*<br>*syringae* (bacterial<br>speck) | Lorrain et al.,<br>2004; Bouchez et<br>al., 2007 |
| **WRKY25**<br>(AT2G30250*) | WRKY<br>DNA-binding<br>protein 25 | Defense suppression<br>(SA dependent) | *Pseudomonas syringae* (bacterial speck) | Zheng et al., 2007 |

Sources:
\* TAIR database
\*\* Ensembl Plants
\*\*\* from van Schie and Takken, 2014

**Figure 4.**
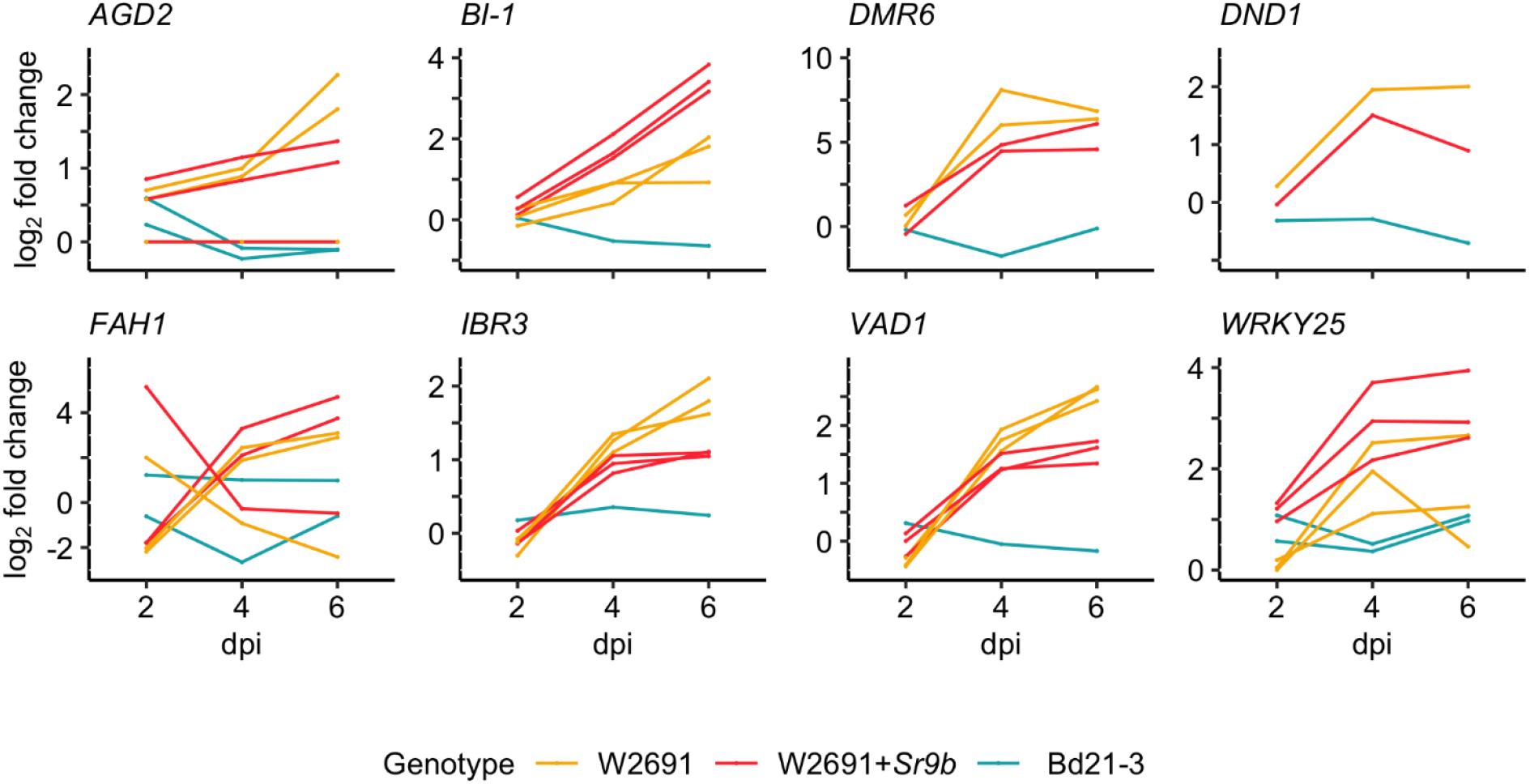
RNAseq expression profile patterns of selected orthogroups containing candidate *S* genes in *T. aestivum* (W2691 and W2691+*Sr9b*) and *B. distachyon* (Bd21-3) genotypes throughout infection with *P. graminis* f. sp. *tritici*. Log2 fold change values for all gene orthologs are presented for each infected genotype compared to the mock treatment per sampling time point. Gene IDs, average FPKM values, orthogroup, and co-expression cluster identifiers are presented in Table S7.

The phylogenetic relationships of the orthogroups to known *S* genes were confirmed using NGphylogeny (**Figure S2**). A phylogenetic tree for *DND1* was not generated since the orthogroup (OG0018857) only contains three genes (TraesCS5D01G404600, BdiBd21-3.1G0110600, and AT5G15410). Complete sets of *T. aestivum* homeologs from the three subgenomes were found in four out of the eight examined orthogroups. There were only two of three expected *T. aestivum* homeologs in the *DMR6* orthogroup, with TraesCS4B02G346900 and TraesCS4D02G341800 representing the B and D subgenomes, respectively. A tblastn of these sequences to chromosome 4A revealed TraesCS4A02G319100, a partial match of 30-31% identity (1e-42 to 1e-44). This gene has low expression and is found in orthogroup OG0006808, which contains two other *T. aestivum* genes, one *B. distachyon* gene, two *A. thaliana* genes, and one *H. vulgare* gene (**Tables S4 and S6**). Despite the low sequence similarity, TraesCS4A02G319100 and TraesCS4B02G346900 are at more similar positions (4A:608043459 and 4B:640532917, respectively) to each other than to TraesCS4D02G341800 (4D:498572979). A tblastn to the entire genome revealed 20 other matches for the two *DMR6* orthologs with 31-74% identity. Thus, it does not seem that the *T. aestivum* genome reference (Chinese Spring) contains a homeolog of *DMR6* in the A genome.

Another *S* gene orthogroup without full subgenome representation was OG0018857 which contained *DND1*. This orthogroup only has one *T. aestivum* gene, TraesCS5D02G404600 from subgenome D. A tblastn to chromosomes 5A and 5B resulted in matches with high identity on both 5A (TraesCS5A02G395300, 94%, 6e-159) and 5B (89%, 1e-176). TraesCS5A02G395300 is present in orthogroup OG0048986 as a singleton with low expression in W2691 (FPKM = 2.61) and W2691+*Sr9b* (FPKM = 1.65), and TraesCS5B02G400100 is included in orthogroup OG0048844 as a singleton as well with notable expression at 6 dpi in infected W2691 (FPKM = 4.28) and low expression in W2691+*Sr9b* (FPKM = 1.19) (**Table S5 and S6**). A tblastn to the entire genome identified 19 other candidates with identity 33-97%. The most notable matches with high identity were TraesCS7B02G161600 (97%, 4e-152) and TraesCS3B02G306700 (97%, 2e-147), which are the only two genes together in orthogroup OG0027858. Both top matches had essentially no expression in either *T. aestivum* genotype (FPKM = 0 to 0.07) (**Table S6**). A third genomic region on chromosome 2B also has 97% identity, but is annotated as a nested repeat.

For *FAH1*, three *T. aestivum* orthologs are present in the orthogroup OG0006155, but one is from subgenome A (TraesCS5A02G019200) while the other two are from subgenome D (TraesCS5D02G024600 and TraesCS5D02G424200). A tblastn of all three sequences to chromosome 5B revealed two matches, TraesCS5B02G016700 (87%, 3e-49) and TraesCS5B02G418800 (62/87%, 2e-64/2e-49), while a tblastn to the entire genome uncovered a partial match on 5A (TraesCS5A02G416500, 46-62%, 7e-68-1e-104) and a partial match on 3D which was not annotated (68-89%, 6e-34-3e-43). All three annotated genes are in singleton orthogroups (TraesCS5B02G016700, OG0056228; TraesCS5B02G418800, OG0055841; TraesCS5A02G416500, OG0057903) and have low expression in both W2691 and W2691+*Sr9b* (FPKM = 0.05 to 1.8) (**Table S5 and S6**). Orthogroup OG0005265 for *AGD2* is similar to the orthorgoup for *FAH1*, having one A subgenome representative (TraesCS4A02G116000) and two D subgenome representatives (TraesCS4D02G189600 and TraesCS7D02G452900). The tblastn of these sequences to chromosome 4B revealed one possible match with two annotations in the same location (61-62%, 9e-108-1e-113), TraesCS4B02G264500 on the - strand and TraesCS4B02G264400 on the + strand. The former is a singleton in orthogroup OG0047603 with low expression in W2691 (FPKM = 0.09) and high expression in infected W2691+*Sr9b* at 6 dpi (FPKM = 4.64), while the latter is in OG0015484 with several other genes and is not highly expressed in either *T. aestivum* genotype (FPKM = 1.3 to 1.7) (**Table S6**). The tblastn to the entire genome revealed several hits of identity varying between 22% and 97%.

### 3.4 Gene co-expression network analysis

To further explore potential processes and novel genes linked to stem rust susceptibility, a gene co-expression network for *B. distachyon* and each *T. aestivum* genotype using the mock and infected RNA-seq data at each timepoint was constructed (**Table S8**). The complete Bd21-3 network has 572,179 edges that connected 21,746 nodes (55.7% of protein-coding genes), while the W2691 and W2691+*Sr9b* networks are larger (W2691: 3,433,279 edges, 49,082 nodes, 45.6% of protein-coding genes; W2691+*Sr9b*: 3,817,404 edges, 49,000 nodes, 45.5% of protein-coding genes). The *B. distachyon* network was expected to be smaller as it represents a diploid species with fewer annotated genes (39,068), while the hexaploid wheat contains more gene annotations (107,891). There are 189 clusters with more than 10 genes in Bd21-3, 258 in W2691, and 391 in W2691+*Sr9b*. Thus, more genes have similar expression patterns in W2691+*Sr9b* than in W2691, and Bd21-3 has the lowest number of genes with similar patterns. The eight *S* gene orthogroups of interest are represented by 14 clusters in W2691 (cluster IDs: 0, 3, 4, 5, 8, 11, 13, 60, 110, 178, 11114, 11235, 12377, 20128), 11 in W2691+*Sr9b* (cluster IDs: 0, 2, 4, 112, 1139, 1916, 2729, 2772, 3133, 10229, 11079), and 11 in Bd21-3 (cluster IDs: 3, 4, 35, 51, 272, 513, 652, 1359, 1662, 1848, 3087) (**Table S9**). Some orthogroups are represented across multiple clusters, while others are only represented in singleton clusters. The ortholog clusters in *B. distachyon* contain fewer genes than the corresponding W2691 and W2691+*Sr9b* ortholog clusters.

GO enrichment tests using GOslim annotations were conducted on the clusters to investigate functional processes. Across all eight *S* gene orthogroups, at least one gene from each is in a cluster with GO enrichment in at least one genotype (**Figure 5**). *DMR6, FAH1*, and *WRKY25* are the only candidates to have enrichment in all three genotypes, *AGD2* and *DND1* only has enrichment in W2691, and *BI-1, IBR3*, and *VAD1* have enrichment in both W2691 and W2691+Sr9b. Terms commonly enriched in the *T. aestivum* genotypes include the Golgi apparatus (GO:0005794), endosome (GO:0005768), endoplasmic reticulum (GO:0005783), protein binding (GO:0005515), transporter activity (GO:0005215), vacuole (GO:0005773), and peroxisome (GO:0005777) (**Figure 5**). Only one GO term, catalytic activity (GO:0003824) is unique to Bd21-3, with other terms like DNA-binding transcription factor activity (GO:0003700) being enriched in the Bd21-3 and *T. aestivum* genotypes.

**Figure 5.**
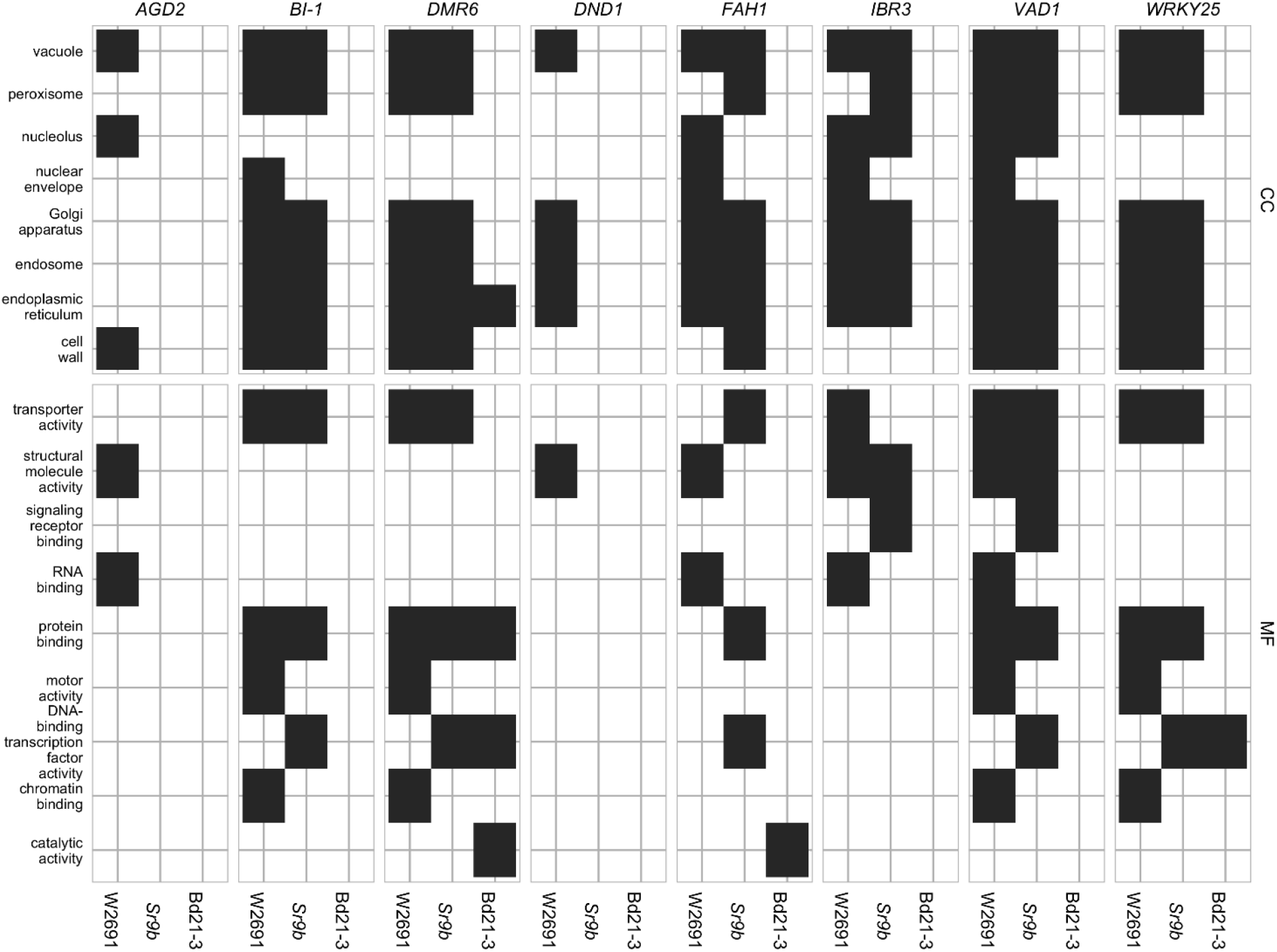
GOslim term enrichment for all genes in co-expression gene clusters containing *S* gene orthologs in *T. aestivum* and *B. distachyon*. The y-axis shows GOslim terms separated into categories: cellular component (CC) and molecular function (MF).

For each genotype a cluster containing one or more orthologs of *DND1, VAD1*, and *DMR6* was selected as examples for presentation (**Figure 6**). Selection criteria for these examples included 1) higher expression in infected than in mock treatments in *T. aestivum* and 2) varied cluster sizes across genotypes. *DND1* is represented by TraesCS5D02G404600 within cluster 4 in the W2691 genotype (557 genes), by TraesCS5D02G404600 within cluster 122 in the W2691+*Sr9b* genotype (21 genes) and by BdiBd21-3.1G0110600 within cluster 652 in the Bd21-3 genotype (4 genes) (**Figure 6A, Table S8**). *VAD1* represents a mid-point between *DND1* and *DMR6*, with the large cluster 0 (TraesCS2D02G236800) representing VAD1 for the W2691 genotype (4527 genes), a singleton cluster (cluster 3087) for the Bd21-3 genotype (1 gene, BdiBd21-3.1G0357000), and the large cluster 0 (TraesCS2D02G236800) for the W2691+*Sr9b* genotype (3400 genes) (**Figure 6B, Table S8**). *DMR6* is also represented by cluster 0 (TraesCS4B02G346900 and TraesCS4D02G341800) for both W2691 and W2691+*Sr9b*; however, cluster 4 representing *DMR6* in Bd21-3 (BdiBd21-3.1G1026800) is larger than in the previous examples (443 genes) (**Figure 6C, Table S8**). In all cases, the *S* gene candidates are not the most differentially-expressed genes at 6 dpi among the *T. aestivum* genotype clusters; the most differentially expressed gene at 6 dpi in cluster 0 is TraesCS7A02G157400 (not functionally annotated) in W2691 (log2FC = 13.64) and TraesCS1A02G266000 (IPR002921:Fungal lipase-like domain IPR029058:Alpha/Beta hydrolase fold IPR033556:Phospholipase A1-II) in W2691+*Sr9b* (log2FC = 14.61). For cluster 4 in W2691, TraesCS4D02G120200 (IPR001471:AP2/ERF domain IPR016177:DNA-binding domain superfamily IPR036955:AP2/ERF domain superfamily) is the most differentially expressed gene at 6 dpi (log2FC = 12.71), while for cluster 122 in W2691+*Sr9b* it is TraesCS1A02G276800 (IPR013087:Zinc finger C2H2-type IPR036236:Zinc finger C2H2 superfamily) (log2FC = 3.83). In *B. distachyon*, BdiBd21-3.1G0110600, which is the Bd21-3 ortholog to *A. thaliana DND1*, is most differentially expressed in the cluster representing *DND1* and was highly downregulated in infected tissue at 6 dpi (log2FC = −0.70). By necessity the most differentially expressed gene in the network representing *VAD1* in *B. distachyon* is the ortholog of *VAD1*, as Bd21-3 cluster 3087 is a singleton cluster. The most differentially expressed Bd21-3 gene in cluster 4 representing *DMR6* is BdiBd21-3.2G0466100 (log2FC = 0.28). This gene is annotated as a Leucine-rich repeat protein kinase family protein due to homology with the *A. thaliana* gene AT1G79620, though the orthology analysis places these genes in different clusters (OG0019394 and OG0010938, respectively). All clusters representing the eight *S* gene candidates are shown in **Figure S3**.

**Figure 6.**
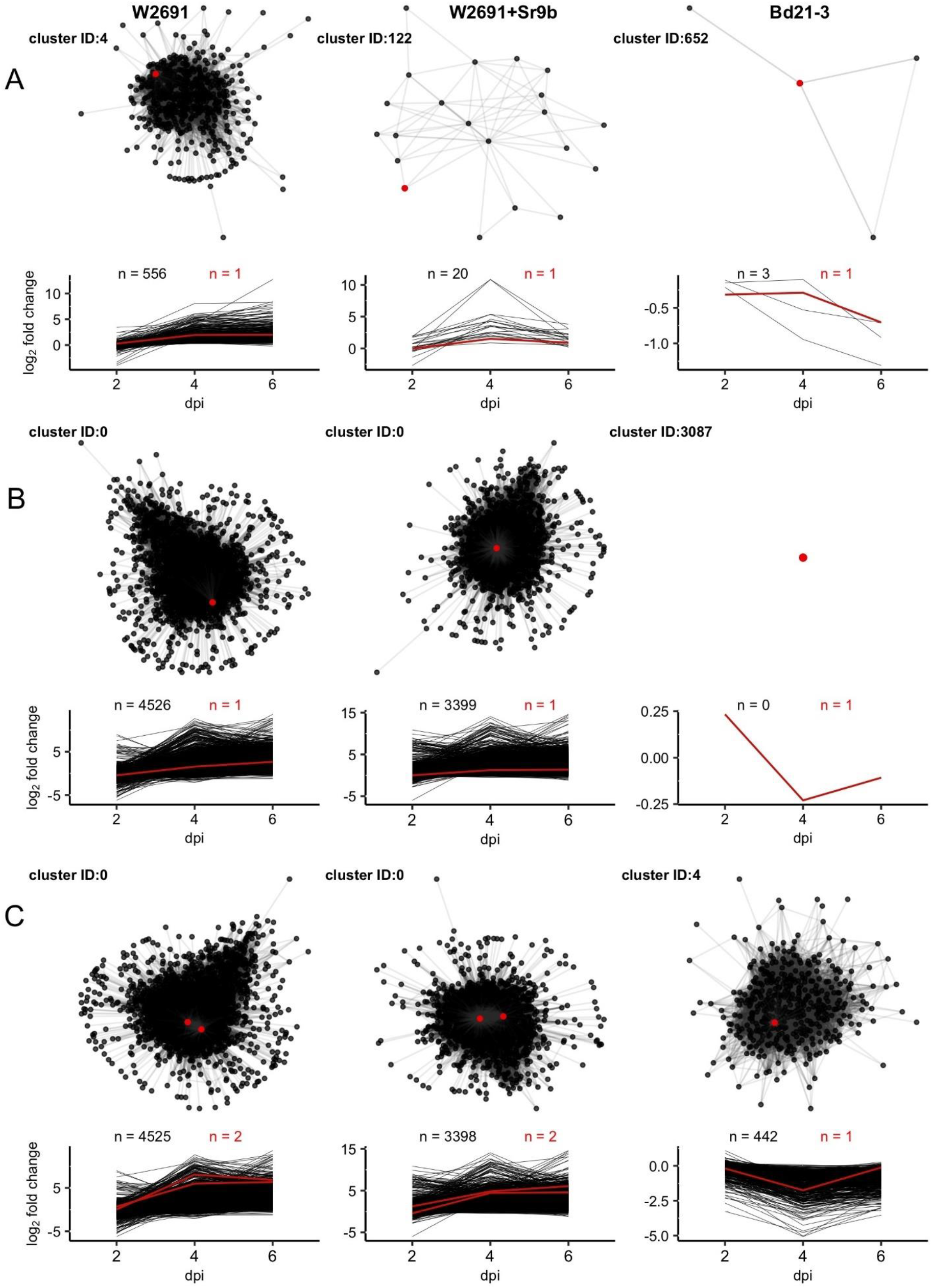
Network diagrams for clusters containing orthologs of **(A)** *DND1*, **(B)** *VAD1*, and (**C**) *DMR6* with corresponding plots showing log2 fold change of all nodes across 2, 4, and 6 dpi. Only connections with Z >= 3 are shown. Red lines, points, and counts represent *T. aestivum* and *B. distachyon* orthologs of *S* genes. Cluster identifiers (IDs) and gene names presented, left to right: *DND1:* 4 (TraesCS5D02G404600), 122 (TraesCS5D02G404600), 652 (BdiBd21-3.1G0110600); *VAD1*: 0 (TraesCS2D02G236800), 0 (TraesCS2D02G236800), 3085 (BdiBd21-3.1G0357000); *DMR6*: 0 (TraesCS4B02G346900), 0 (TraesCS4D02G341800), 4 (BdiBd21-3.1G1026800).

## 4 Discussion

Susceptibility (*S*) genes are an essential component of compatible plant pathogen interactions (Engelhardt et al., 2018). The opportunity to genetically manipulate such genes to engineer disease resistance in important crops such as *T. aestivum* has captured significant scientific interest in recent years. However, our understanding of the genetic basis of disease susceptibility in cereals is limited to a few examples (van Schie and Takken, 2014; Engelhardt et al., 2018). Thus, important questions regarding the biological functions of these genes and their activation remain to be answered. As a first step to uncover putative stem rust *S* genes, we conducted a comparative RNA-seq experiment coupled with gene co-expression network analysis to determine transcriptional responses in *T. aestivum* genotypes and *B. distachyon* Bd21-3. We compared a compatible interaction (W2691) with an incompatible interaction controlled by the race-specific resistance gene *Sr9b* in the same genetic background (W2691*+Sr9b*). *Sr9b* restricts pathogen growth; however, it also allows the development of small sporulating colonies of a *Pgt* isolate which belongs to the race SCCL (Zambino et al., 2000). A more stringent incompatibility scenario is given by Bd21-3 genotype of *B. distachyon*, which allows restricted colony formation of *Pgt* without sporulation. These observations were consistent with previous descriptions of *B. distachyon* as a non-host to rust pathogens (Figueroa et al., 2013, 2015, Omidvar et al., 2018). Thus, a strength of this study is the survey of molecular responses associated with increasing levels of susceptibility.

Consistent with findings from other transcriptomic studies of wheat-rust interactions (Dobon et al., 2016; Manickavelu et al., 2010; Chandra et al., 2016; Zhang et al., 2014; Yadav et al., 2016), major transcriptional changes were detected in response to infection in both *T. aestivum* and *B. distachyon*, which reflect the complexity of these plant-microbe interactions. A significantly higher number of up- or down-regulated genes were found in *T. aestivum* than *B. distachyon*. The greater fungal colonization of *T. aestivum* as indicated by *in planta* fungal growth assays of *Pgt* is likely a result of the pathogen’s failure to effectively manipulate the metabolism of *B. distachyon*. GOslim term analyses indicated an enrichment for Golgi apparatus, peroxisome, vacuole, and cell wall related functions in up-regulated genes in *T. aestivum*. These results are not surprising as a large proportion of immune receptors and plant defense signaling components play a role in plant-microbe interactions (Couto and Zipfel, 2016; Dodds and Rathjen, 2010). The plant Golgi apparatus and peroxisomes have been reported as targets of effectors from various pathogenic filamentous fungi (Robin et al., 2018). The enrichment of these GO terms in up-regulated genes in *T. aestivum* suggests that these cellular components may be direct or indirect targets for effectors derived from *Pgt*. Analyses with the full GO term set revealed many enriched terms among downregulated genes related to photosynthesis in W2691 and W2691+*Sr9b*; a decrease in chlorophyll and photosynthetic activity has been previously reported in wheat infected with *Pgt* (Berghaus and Reisener, 1984; Moerschbacher et al., 1994).

Several *S* genes to diverse pathogens have been identified or postulated in various plant species (van Schie and Takken, 2014; Engelhardt et al., 2018). While this area of research for cereal rust pathogens is in its infancy, positive results from other pathosystems make a strong case to consider the modification of *S* genes as an approach to deliver durable and broad-spectrum disease resistance. So far, only a few host-delivered avirulence effectors, *AvrSr50* (Chen et al., 2017), *AvrSr35* (Salcedo et al., 2017), *AvrSr27* (Upadhyaya et al., accepted) from any cereal rust fungi have been isolated. These were identified in *Pgt* and how these effectors disrupt defense responses in compatible interactions remains unknown. Future research seeking to identify which plant proteins these effectors target will help elucidating *S* genes or processes required for stem rust susceptibility.

Here, expression patterns of gene orthologs in *T. aestivum* and *B. distachyon* corresponding to previously characterized *S* genes in *H. vulgare* and *A. thaliana* were examined to develop a framework to study *S* genes in wheat. A key focus of this study was to develop a workflow to extract orthologs with high expression in stem rust susceptible *T. aestivum*, but low expression in either *T. aestivum* with intermediate resistance, or *B. distachyon*. To link these candidate *S* genes with the biological pathways in *T. aestivum* and *B. distachyon*, we constructed gene co-expression networks, which can be explored to determine the role of components of these pathways and the complex interplay towards regulation of susceptibility in *Pgt*-*T. aestivum* interactions.

The biological functions of *S* genes in compatible-plant microbe interactions are diverse, as these genes play roles in a wide array of events that are critical for pathogen accommodation and survival (Engelhardt et al., 2018). Some of these susceptibility genes can act as negative regulators of immune responses, such as PTI, cell death, and phytohormone-related defense. Our study determined that *T. aestivum* orthologs of the BAX inhibitor-1 (*BI-1*) gene in *H. vulgare* are candidate *S* genes, as these were upregulated in W2691 (6 dpi) and W2691+*Sr9b* (4-6 dpi) whereas their expression in *B. distachyon* was not affected. *BI-1* is an endoplasmic reticulum membrane-localized cell death suppressor in *A. thaliana*, and its wheat ortholog *TaBI-1* (accession GR305011) is proposed to contribute to susceptibility in *T. aestivum* to the biotrophic pathogen *Puccinia striiformis* f. sp. *tritici* (Wang et al., 2012). Interestingly, the highest upregulation of the *BI-1* was detected in the W2691+*Sr9b* genotype where it is necessary to regulate a HR upon *Pgt* recognition. Given this result it should be examined if *BI-1* may be a conserved plant *S* factor to wheat rust fungi. Various orthologs of *FAH1*, which encodes a ferulate *5-*hydroxylase in *A. thaliana*, were upregulated in the *T. aestivum* genotypes upon *Pgt* infection (Mitchell and Martin, 1997). According to studies in *A. thaliana* FAH1 plays a role in *BI-1*-mediated cell death suppression through interaction with cytochrome b5 and biosynthesis of very-long-chain fatty acids (Nagano et al., 2012). Additional findings further suggest that *Pgt* can also interfere with cell death signaling by altering *VAD1* expression. The *VAD1* gene encodes a putative membrane-associated protein with lipid binding properties and it is proposed to act as negative regulator of cell death (Lorrain et al., 2004; Khafif et al., 2017). Transcriptional activation of *VAD1* has been shown to occur in advanced stages in plant pathogen interactions (Bouchez et al., 2007). We detected an upregulation of *VAD1* orthologs in *T. aestivum* at 6 dpi, which is considered a late infection stage in the establishment of rust colonies.

Salicylic acid (SA) is a key phytohormone required to orchestrate responses to many pathogens (Ding and Ding, 2020). Similar to VAD1 whose function as a *S* factor is SA-dependent, we also uncovered other upregulated genes that may also participate in defense suppression. The orthologs of the *DMR6* are highly upregulated in *T. aestivum* at 4 and 6 dpi in both compatible and incompatible interactions. As characterized in *A. thaliana, DMR6* encodes a putative 2OG-Fe(II) oxygenase that is defense-associated and required for susceptibility to downy mildew through regulation of the SA pathway (Van Damme et al., 2008, Zhang et al., 2017). The role of *DMR6* in disease susceptibility holds significant promise to control diverse pathogens. For instance, mutations in *DMR6* confer resistance to hemibiotrophic pathogens *Pseudomonas syringae* and *Phytophthora capsici* (Zeilmaker et al., 2014) and silencing of *DMR6* in potato increases resistance to the potato blight causal agent, *P. infestans* (Sun et al., 2016). It has also been shown that the *H. vulgare* ortholog genetically complements *DMR6* knock-out *A. thaliana* lines and restores susceptibility to *Fusarium graminearum* (Low et al., 2020). Gene orthologs of *DND1* were also identified as upregulated in both *T. aestivum* genotypes. The gene *DND1* encodes a cyclic nucleotide-gated ion channel and its activity is also related to SA regulation (Clough et al., 2000). Mutations in *A. thaliana DND1* display enhanced resistance to viruses, bacteria and fungal pathogens (Genger et al., 2008; Jurkowski et al., 2004; Yu et al., 2000; Sun et al., 2017). We also noted that several wheat orthologs of the *A. thaliana* gene *IBR3* also increased in expression as *Pgt* infection advanced. The role of *IBR3* in susceptibility to *P. syringae* in *A. thaliana* has been confirmed by mutations and overexpression approaches (Huang et al., 2013). Consistent with our results, *IBR3* is upregulated in *A. thaliana* upon infection by *P. syringae*.

Plant transcriptional reprogramming triggered by pathogen perception is often mediated by WRKY transcription factors through activation of the MAP kinase pathways (Eulgen and Somssich, 2007; Rushton et al., 2010). Here, we detected an upregulation of the expression of *WRKY25* orthologs that was most prominent at 6 dpi in the W2691+*Sr9b* genotype. The Arabidopsis gene *AtWRKY25* is induced in response to the bacterial pathogen *Pseudomonas syringae* and the SA-dependent activity of *AtWRKY25* is also linked to defense suppression (Zheng et al., 2007). According to results from this study, the contribution of orthologs of *AGD2* to stem rust susceptibility in *T. aestivum* should also be examined. *AGD2* encodes an aminotransferase and participates in lysine biosynthesis at the chloroplast (Song et al., 2004). Given that several oomycete and fungal effectors target the chloroplast (Kretschmer et al., 2020), effector research in cereal rust pathogens will be crucial to determine if these pathogens also target this organelle.

A classic example of *S* genes in barley is given by the *Mlo* gene (Jørgensen, 1992) in which a recessive mutation results in broad spectrum resistance to *Blumeria graminis* f. sp. *hordei*, the causal agent of powdery mildew. The *Mlo* gene family is highly conserved across monocot and dicot plants and gene editing of *Mlo* homeologs in wheat confers resistance to powdery mildew (Acevedo-Garcia et al., 2017). Interestingly, *Mlo* genes in *T. aestivum* have not been reported to provide protection against cereal rust diseases. Consistent with this, this study did not detect a significant change in the expression of *Mlo* alleles in *T. aestivum* genotypes (W2691 and W2691+*Sr9b*) over the course of the experiment.

One caveat of this study is that some *S* genes in *T. aestivum* for *Pgt* may not be found in model species like *A. thaliana* or detected using other pathogens. However, this is a first step to identify candidates to guide functional studies. While in this study we focused on orthologous *S* genes, the gene co-expression networks presented here are excellent resources to identify additional candidate *S* factors. It is possible that some of the genes included in clusters of these networks are part of the regulatory process that control expression of *S* genes or are part of essential pathways although their function may not be characterized yet in other systems. Future functional studies are required to validate the function of these genes in *T. aestivum* as *S* factors for rust infection and determine if these can be exploited for agricultural practice. A key aspect for the success of these novel approaches is the absence of plant developmental defects resulting from mutations of *S* genes. In some cases, the loss-of-function of negative regulators leads to constitutive activation of plant defense responses that manifest as poor growth or lesion-mimic phenotypes among other pleiotropic effects (Büschges et al., 1997; Ge et al., 2016). VIGS-mediated transient gene silencing (Lee et al., 2015), RNAi-mediated silencing (Sun et al., 2016; Helliwell and Waterhouse, 2003; Waterhouse and Helliwell, 2003), TILLING populations include some of the approaches to explore the potential use of these *S* gene candidates. Gene editing technologies through Zinc Finger nucleases, TALENs, CRISPR/Cas9 systems also offer options to generate transgene free plants (Urnov et al., 2010; Gaj et al., 2013; Luo et al., 2019; Kim et al., 2017; Jia et al., 2017; Wang et al., 2014; Jia et al., 2017; Nekrasov et al., 2017). In conclusion, as the demands for multi-pathogen durable disease resistance rise, our ability to target *S* genes may serve as a sound approach to harness genetic diversity and maximize the resources to meet critical these grand challenges.

## 6 Author Contributions

MF, CDH, CLM, and SFK conceived and designed the study; VO, MEM, FL, and MF conducted the experiments; ECH, EG, JMM, RDC, CDH, SPG, JPV, and MEM contributed to data analysis. ECH, BJS, SFK, CDH and MF interpreted results. ECH, VO, CDH, and MF wrote the manuscript; all authors contributed to manuscript editing, revisions and approved the submitted version.

## 8 Acknowledgments

We thank staff at the Minnesota Supercomputing Institute at the University of Minnesota for technical assistance.

## 10 Data Availability Statement

Sequence data was deposited in NCBI under BioProject PRJNA483957 (**Table S1**). Unless specified otherwise, scripts and files for analysis and visualizations are available at https://github.com/henni164/stem_rust_susceptibility.

---

**Supplementary Figure 1.**
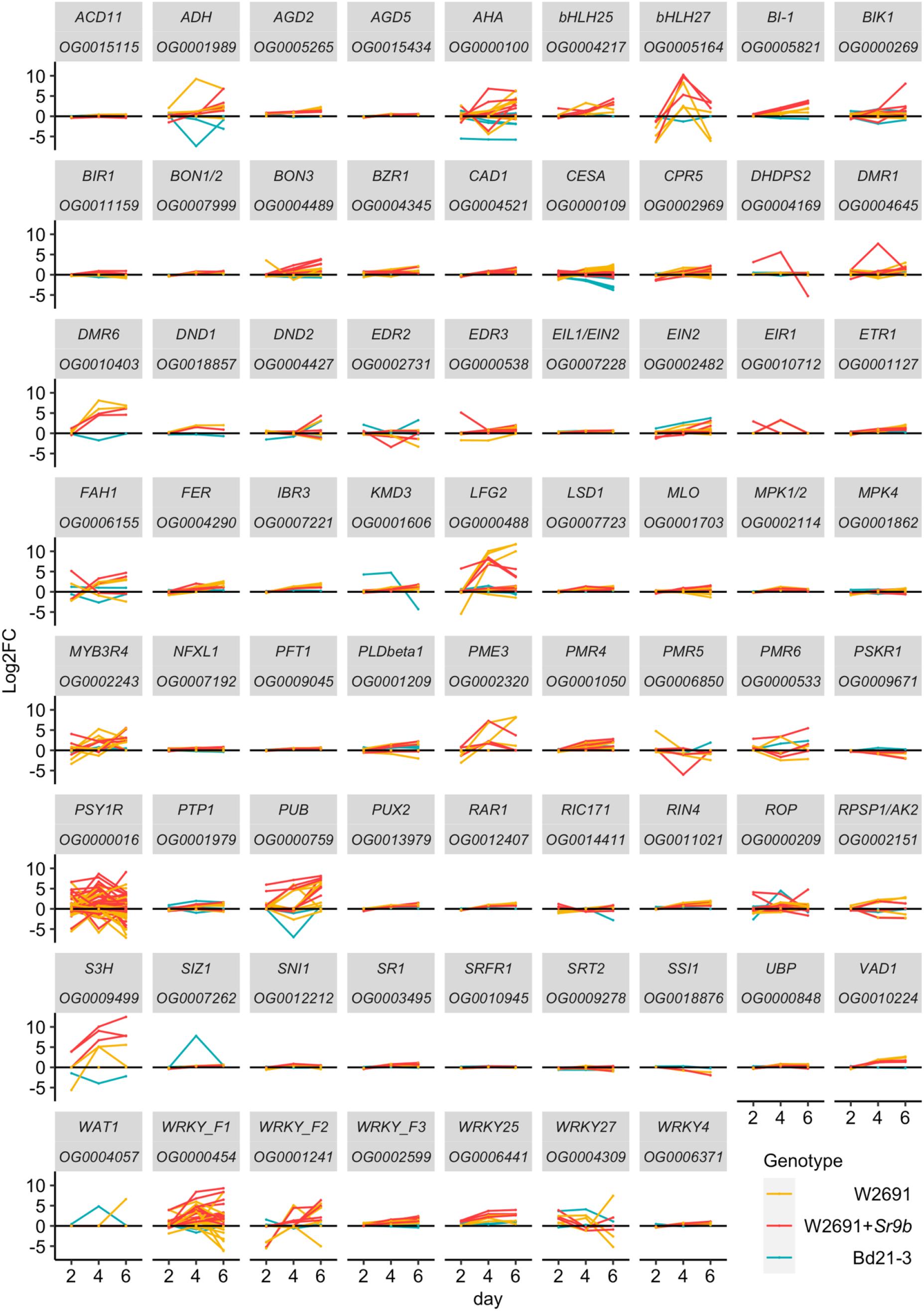
Expression profile patterns of orthogroups containing candidate *S* genes in *T. aestivum* (W2691 and W2691+*Sr9b*) and *B. distachyon* (Bd21-3) genotypes throughout infection with *P. graminis* f. sp. *tritici*. Log2 fold change values (y-axis) for all gene orthologs are presented per sampling time point (x-axis). Name of *S* gene and orthogroup identifier are shown in each graph.

**Supplementary Figure 2.**
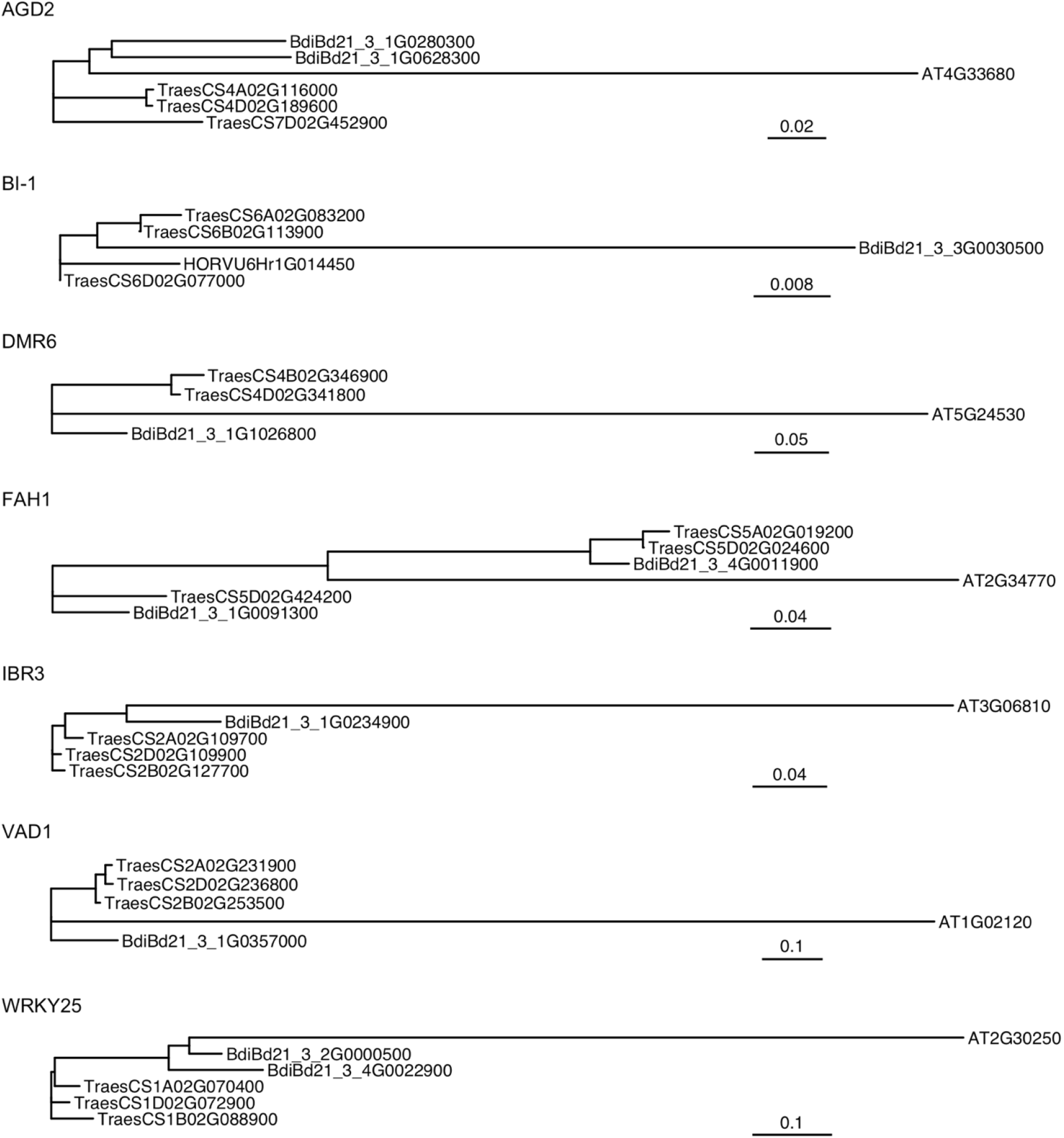
Molecular phylogenetic analysis of amino acid sequences of orthologous genes for five of the six S genes of interest. The orthogroup for DND1 only included three genes (One *A. thaliana* susceptibility gene, one *T. aestivum* ortholog, and one *B. distachyon* ortholog) and so no phylogenetic tree was generated. Scale bars represent nucleotide substitutions per site.

**Supplementary Figure 3.**
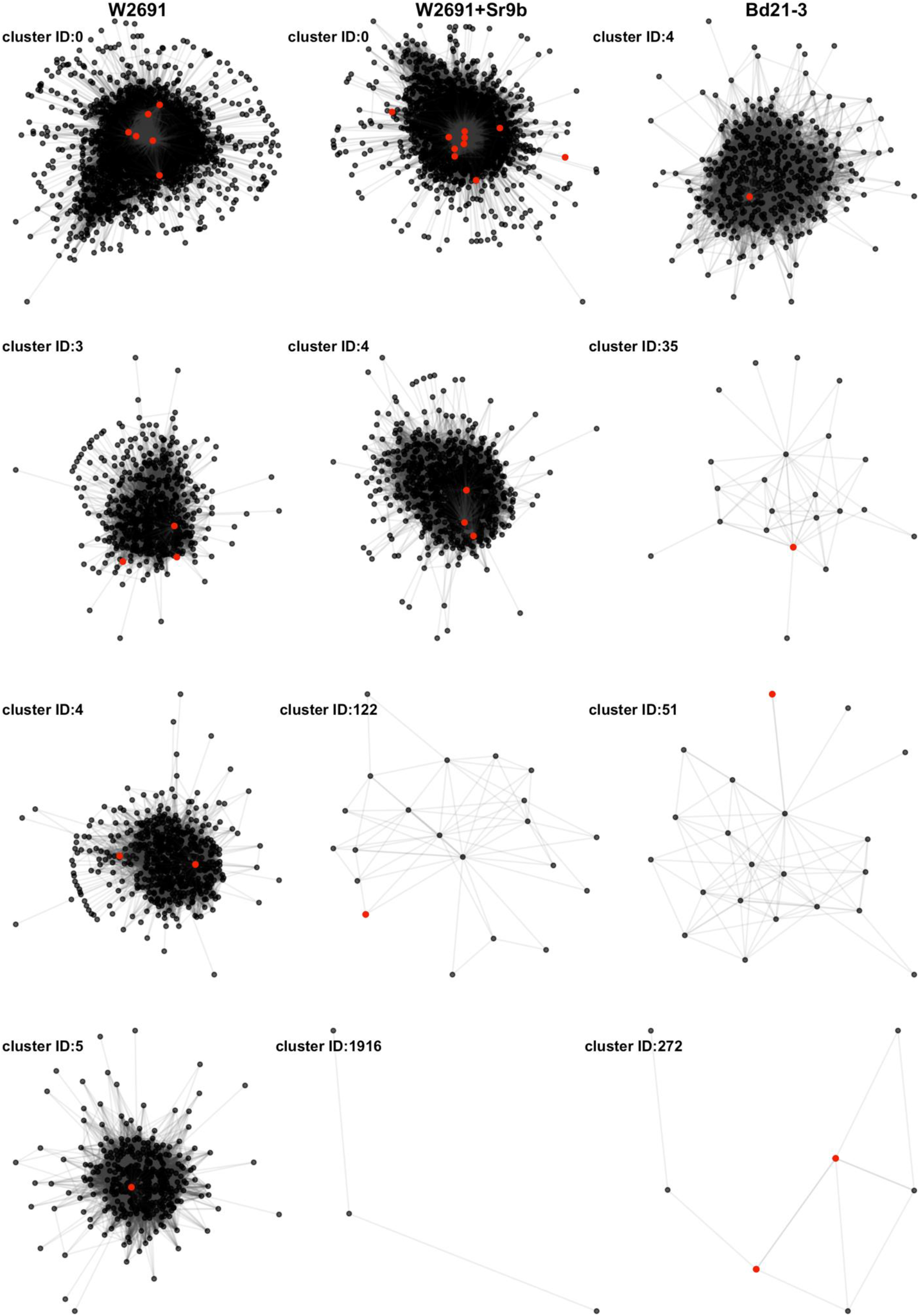
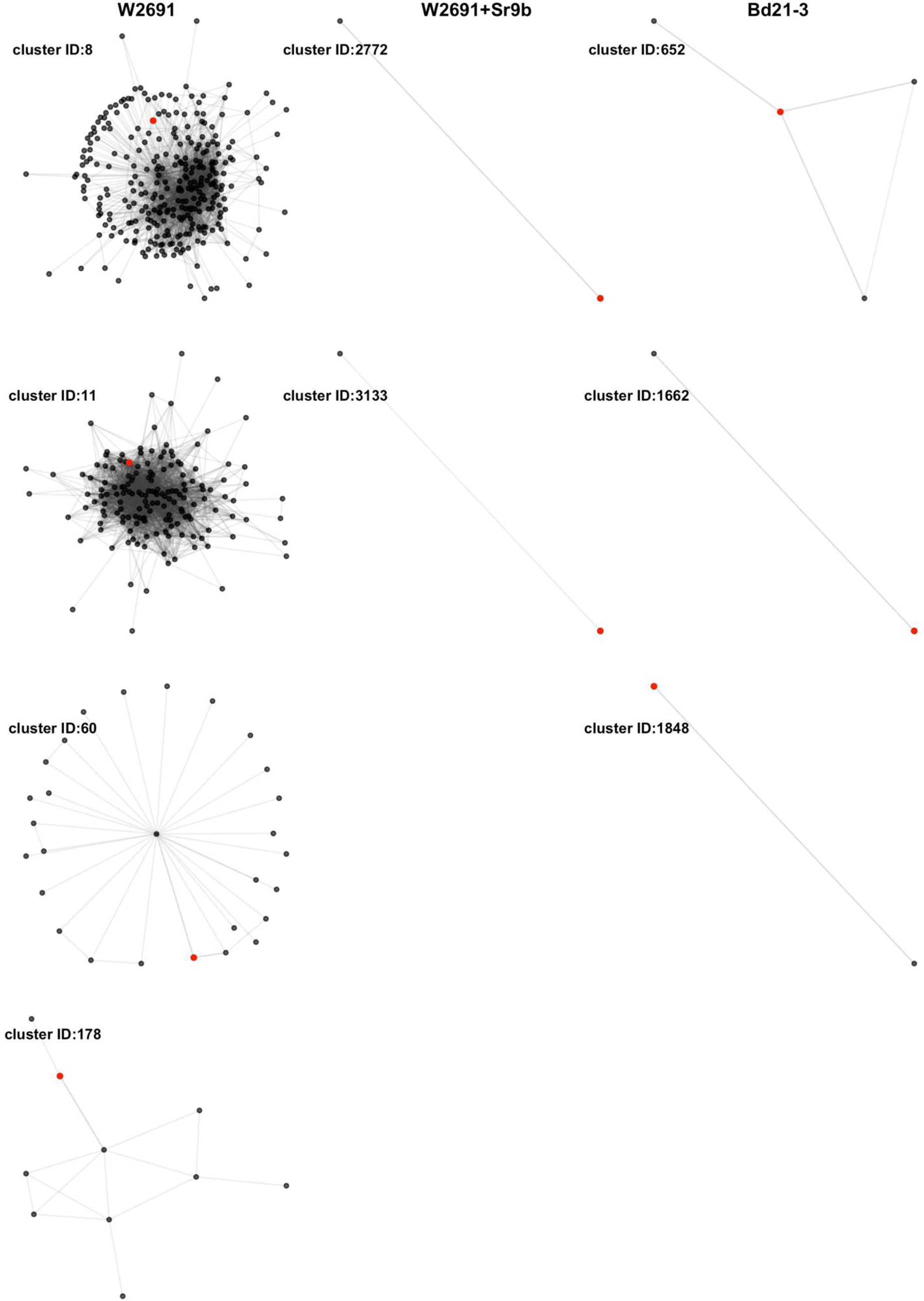
All clusters (nodes > 1) containing a *T. aestivum* or *B. distachyon* ortholog of the eight susceptibility candidates. Red points represent *T. aestivum* and *B. distachyon* orthologs of *S* genes. Gene IDs of members in co-expression clusters are presented in Table S9.

## Supplementary Tables

**Table S1**. RNA-seq reads, NCBI accession numbers and mapping statistics (excel file).

**Table S2**. Enriched GO terms (Observed count over expected count) among up- and down-regulated genes at three time points across the two *T. aestivum* genotypes and *B. distachyon* Bd21-3.

**Table S3**. List of candidate susceptibility genes and orthogroups (excel file).

**Table S4**. List of orthogroups containing two or more genes including gene IDs (excel file).

**Table S5**. List of singleton orthogroups and gene IDs (excel file).

**Table S6**. Average gene expression (FPKM), orthogroup, GO terms, and cluster numbers associated with all genes in W2691, W2691+*Sr9b*, Bd21-3 (excel file).

**Table S7**. Average gene expression (FPKM), orthogroup, GO terms, and cluster numbers associated with eight *S* gene candidates in W2691, W2691+*Sr9b*, Bd21-3 (excel file).

**Table S8**. Co-expression network data for W2691, W2691+*Sr9b*, Bd21-3 genotypes using the mock and infected RNA-seq data (text file).

Available at https://github.com/henni164/stem_rust_susceptibility

**Table S9**. Co-expression network data for clusters containing *S* gene candidates (text file). Available at https://github.com/henni164/stem_rust_susceptibility

## References

Ahn IP. (2007). Disturbance of the Ca(2+)/calmodulin-dependent signaling pathway is responsible for the resistance of Arabidopsis dnd1 against Pectobacterium carotovorum infection. Mol. Plant Pathol. 8, 747–59

Alaux, M., Rogers, J., Letellier, T., Flores, R., Alfama, F., Pommier, C., et al. (2018). Linking the International Wheat Genome Sequencing Consortium bread wheat reference genome sequence to wheat genetic and phenomic data. Genome Biol. 19, 111

Alexa, A. and Rahnenfuhrer, J. (2020). topGO: Enrichment Analysis for Gene Ontology. R package version 2.40.0.

Anders, S., Pyl, P.T., Huber, W. (2015). HTSeq—a Python framework to work with high-throughput sequencing data. Bioinformatics. 31, 166–169

Ayliffe, M., Singh, D., Park, R., Moscou, M., and Pryor, T. (2013). Infection of Brachypodium distachyon with selected grass rust pathogens. Mol. Plant Microbe In. 26, 946–957

Berghaus, R. and Reisener, H.J. (1984). Changes in photosynthesis of wheat plants infected with wheat stem rust (Puccinia graminis f. sp. tritici). Phytopathology. 112, 165–172

Bettgenhaeuser, J., Gardiner, M., Spanner, R., Green, P., Hernández-Pinzón, I., Hubbard, A., et al. (2018). The genetic architecture of colonization resistance in Brachypodium distachyon to nonadapted stripe rust (Puccinia striiformis) isolates. PLoS Genet. 14

Bettgenhaeuser, J., Gilbert, B., Ayliffe, M., and Moscou, M. J. (2014). Nonhost resistance to rust pathogens-a continuation of continua. Front. Plant Sci. 5, 664

Bhattacharya, S. (2017). Deady new wheat disease threatens Europe’s crops. Nature. 542:145–146

Bouchez O., Huard C., Lorrain S., Roby D., Balague C. (2007). Ethylene is one of the key elements for cell death and defense response control in the Arabidopsis lesion mimic mutant vad1. Plant Physiol. 145, 465–77

Bozkurt, T. O., McGrann, G. R., MacCormack, R., Boyd, L. A., and Akkaya, M. S. (2010). Cellular and transcriptional responses of wheat during compatible and incompatible race-specific interactions with Puccinia striiformis f. sp. tritici. Mol. Plant Pathol. 11, 625–640

Briatte, F. (2020). ggnetwork: Geometries to Plot Networks with “ggplot2.”

Büschges, R., Hollricher, K., Panstruga, R. Simons, G., Wolter, M., Frijters, A., et al. (1997). The Barley Mlo Gene: A Novel Control Element of Plant Pathogen Resistance. Cell. 88(5), 695–705

Bushnell, W. R. (1984). 15 - Structural and Physiological Alterations in Susceptible Host Tissue. Pages 477–507 in: The Cereal Rusts W.R. Bushnell and A.P. Roelfs, eds. Academic Press.

Butts, C. T. (2015). network: Classes for Relational Data.

Butts, C. T. (2019). sna: Tools for Social Network Analysis.

Chandra, S., Singh, D., Pathak, J., Kumari, S., Kumar, M., Poddar, R., et al. (2016). De novo assembled wheat transcriptomes delineate differentially expressed host genes in response to leaf rust. PLoS ONE. 11

Chen, B., Jiang, J., Zhou, X. (2007). A TOM1 homologue is required for multiplication of Tobacco mosaic virus in Nicotiana benthamiana. J. Zhejiang Univ. - Sci. B. 8, 256–259

Chen, L.-Q., Hou, B.-H., Lalonde, S., Takanaga, H., Hartung, M. L., Qu, X.-Q., et al. (2010). Sugar transporters for intercellular exchange and nutrition of pathogens. Nature. 468, 527–532

Clough, S. J., Fengler, K. A., Yu, I. C., Lippok, B., Smith, R. K., Jr, and Bent, A. F. (2000). The Arabidopsis dnd1 “defense, no death” gene encodes a mutated cyclic nucleotide-gated ion channel. Proc. Natl. Acad. Sci. USA. 97, 9323–9328

Couto, D., and Zipfel, C. (2016). Regulation of pattern recognition receptor signaling in plants. Nat. Rev. Immunol. 16, 537–552

Ding, P., Ding, Y. (2020). Stories of Salicylic Acid: A Plant Defense Hormone. Trends Plant Sci. 25(6), 549–565

Dobin, A., Davis, C. A., Schlesinger, F., Drenkow, J., Zaleski, C., Jha, S., et al. (2012). STAR: ultrafast universal RNA-seq aligner. Bioinformatics. 29, 15–21

Dobon, A., Bunting, D. C., Cabrera-Quio, L. E., Uauy, C., and Saunders, D. G. (2016). The hostpathogen interaction between wheat and yellow rust induces temporally coordinated waves of gene expression. BMC Genomics. 17, 380

Dodds, P. N., and Rathjen, J. P. (2010). Plant immunity: towards an integrated view of plant-pathogen interactions. Nat. Rev. Genet. 11, 539–548

Duplessis, S., Cuomo, C. A., Lin, Y. C., Aerts, A., Tisserant, E., Veneault-Fourrey, C., et al. (2011). Obligate biotrophy features unraveled by the genomic analysis of rust fungi. Proc. Natl. Acad. Sci. USA. 108, 9166–9171

Eichmann R., Bischof M., Weis C., Shaw J., Lacomme C., et al. (2010). BAX INHIBITOR-1 Is Required for Full Susceptibility of Barley to Powdery Mildew. Mol. Plant Microbe In. 23, 1217–27

Ellis, J. G., Lagudah, E. S., Spielmeyer, W., and Dodds, P. N. 2014. The past, present and future of breeding rust resistant wheat. Front. Plant Sci. 5, 641

Emms, D.M., Kelly, S. (2019). OrthoFinder: phylogenetic orthology inference for comparative genomics. Genome Biol. 20(238)

Engelhardt, S., Stam, R., Hückelhoven, R. (2018). Good Riddance? Breaking Disease Susceptibility in the Era of New Breeding Technologies. Agronomy. 8, 114

Eulgem, T. and Somssich, I. E. (2007). Networks of WRKY transcription factors in defense signaling. Curr. Opin. Plant Bio. 10:366–371

Figueroa, M., Alderman, S., Garvin, D. F., and Pfender, W. F. (2013). Infection of Brachypodium distachyon by formae speciales of Puccinia graminis: early infection events and host-pathogen incompatibility. PLoS ONE. 8

Figueroa, M., Castell-Miller, C. V., Li, F., Hulbert, S. H., and Bradeen, J. M. (2015). Pushing the boundaries of resistance: insights from Brachypodium-rust interactions. Front. in Plant Sci. 6, 558

Flor, H. H. (1971). Current status of the gene-for-gene concept. Annu. Rev. Phytopathology. 9, 275–296

Garnica, D. P., Nemri, A., Upadhyaya, N. M., Rathjen, J. P., and Dodds, P. N. (2014). The ins and outs of rust haustoria. PLoS Pathog. 10

Gaj, T., Gersbach, C.A., Barbas, C.F. (2013). ZFN, TALEN, and CRISPR/Cas-based methods for genome engineering. Trends Biotechnol. 31(7), 397–405

Ge, X.T., Deng, W.W., Lee, Z.Z., Lopez-Ruiz, F.J., Schweizer, P., Ellwood, S.R. (2016). Tempered mlo broad-spectrum resistance to barley powdery mildew in an Ethiopian landrace. Sci. Rep. 7, 29558

Genger, R.K., Jurkowski, G.I., McDowell, J.M., Lu, H., Jung, H.W., Greenberg, J.T., et al. (2008). Signaling pathways that regulate the enhanced disease resistance of Arabidopsis “defense, no death” mutants. Mol. Plant Microbe In. 21, 1285-96

Gilbert, B., Bettgenhaeuser, J., Upadhyaya, N., Soliveres, M., Singh, D., Park, R. F., et al. (2018). Components of Brachypodium distachyon resistance to nonadapted wheat stripe rust pathogens are simply inherited. PLoS Genet. 14

Govrin E.M., Levine A. (2000). The hypersensitive response facilitates plant infection by the necrotrophic pathogen Botrytis cinerea. Curr. Biol. 10, 751–7

Harder, D. E., and Chong, J. (1984). Structure and physiology of haustoria. Pages 416–460 in: The Cereal Rusts, W. Bushnell, ed. Academic Press.

Helliwell, C., Waterhouse, P. (2003). Constructs and methods for high-throughput gene silencing in plants. Methods. 30(4), 289–295

Howe, K. L., Contreras-Moreira, B., De Silva, N., Maslen, G., Akanni, W., Allen, J., et al. (2019). Ensembl Genomes 2020—enabling non-vertebrate genomic research. Nucleic Acids Res. 48, D689–D695

Huang T.Y., Desclos-Theveniau M., Chien C.T., Zimmerli L. (2013). Arabidopsis thaliana transgenics overexpressing IBR3 show enhanced susceptibility to the bacterium Pseudomonas syringae. Plant Biol. (Stuttg) 15, 832–40

Huttenhower C., Hibbs M., Myers C., Troyanskaya O.G. (2006). A scalable method for integration and functional analysis of multiple microarray datasets. Bioinformatics. Dec 1;22(23), 2890–7.

Jia, H.G., Zhang, Y.Z., Orbović, V., Xu, J., White, F.F., Jones, J.B., et al. (2017). Genome editing of the disease susceptibility gene CsLOB1 in citrus confers resistance to citrus canker. Plant Biotechnol. J. 15, 817–823

Jørgensen J.H. (1992). Discovery, characterization and exploitation of Mlo powdery mildew resistance in barley. Euphytica. 63, 141–152.

Jurkowski, G.I., Smith, R.K. Jr., Yu, I., Ham, J.H., Sharma, S.B., Klessig, D.F., et al. (2004). Arabidopsis DND2, a second cyclic nucleotide-gated ion channel gene for which mutation causes the “defense, no death” phenotype. Mol. Plant-Microbe Interact. 17(5), 511–20.

Kellogg, E. A. (2001). Evolutionary history of the grasses. Plant Physiol. 125:1198–1205

Khafif, M., Balagué, C., Huard-Chauveau, C., and Roby, D. (2017). An essential role for the VASt domain of the Arabidopsis VAD1 protein in the regulation of defense and cell death in response to pathogens. PLoS ONE. 12, e0179782

Kim, D., Alptekin, B., Budak, H. (2018). CRISPR/Cas9 genome editing in wheat. Funct. Integr. Genomics. 18, 31–41

Konig S., Feussner K., Schwarz M., Kaever A., Iven T., Landesfeind, M., et al. (2012). Arabidopsis mutants of sphingolipid fatty acid alpha-hydroxylases accumulate ceramides and salicylates. New Phytol. 196, 1086–97

Kretschmer, M., Damoo, D., Djamei, A., Kronstad, J. (2020). Chloroplasts and Plant Immunity: Where Are the Fungal Effectors? Pathogens. 9, 19

Lamesch, P., Berardini, T. Z., Li, D., Swarbreck, D., Wilks, C., Sasidharan, R., et al. (2011). The Arabidopsis Information Resource (TAIR): improved gene annotation and new tools. Nucleic Acids Res. 40, D1202–D1210

Lapin, D., and Van den Ackerveken, G. (2013). Susceptibility to plant disease: more than a failure of host immunity. Trends Plant Sci. 18, 546–554

Lawrence-Dill, C. (2019). GOMAP Wheat Reference Sequences 1.1. 1.1. CyVerse Data Commons. DOI: 10.25739/65kf-jz20

Lee, W. S., Rudd, J. J., Kanyuka, K. (2015). Virus induced gene silencing (VIGS) for functional analysis of wheat genes involved in Zymoseptoria tritici susceptibility and resistance. Fungal Genet. Giol. 79, 84–88

Lemoine, F., Correia, D., Lefort, V., Doppelt-Azeroual, O., Mareuil, F., Cohen-Boulakia, S., et al. (2019). NGPhylogeny.fr: new generation phylogenetic services for non-specialists. Nucleic Acids Res. 47, W260–W265

Lo Presti, L., Lanver, D., Schweizer, G., Reissman, S., and Kahmann, R. (2015). Fungal effectors and plant susceptibility. Annu. Rev. Plant Biol. 66, 513–545

Lorrain S., Lin B., Auriac M.C., Kroj T., Saindrenan P., Nicole, M., et al. (2004). Vascular associated death1, a novel GRAM domain-containing protein, is a regulator of cell death and defense responses in vascular tissues. Plant Cell. 16, 2217–32

Love, M.I., Huber, W., and Anders, S. (2014). Moderated estimation of fold change and dispersion for RNA-seq data with DESeq2. Genome Biol. 15, 550

Low, Y. C., Lawton, M. A., and Di, R. (2020). Validation of barley 2OGO gene as a functional orthologue of Arabidopsis DMR6 gene in Fusarium head blight susceptibility. Sci. Rep-UK. 10, 9935

Luig, N.H. and Watson, I.A. (1972). The role of wild and cultivated grasses in the hybridization of formae speciales of Puccinia graminis. Aust. J. Biol. Sci. 25(2), 335–342

Luo, M., Li, H., Chakraborty, S., Morbitzer, R., Rinaldo, A., Upadhyaya, N., et al. (2019). Efficient TALEN-mediated gene editing in wheat. Plant Biotechnol. J. 17, 2026–2028.

Manickavelu, A., Kawaura, K., Oishi, K., Shin-I, T., Kohara, Y., Yahiaoui, N., (2010). Comparative gene expression analysis of susceptible and resistant near-isogenic lines in common wheat infected by Puccinia triticina. DNA Res. 17(4), 211–222

Martin, M. (2011). Cutadapt removes adapter sequences from high-throughput sequencing reads. EMBnet.journal, 17(1), 10-12. doi.org/10.14806/ej.17.1.200

McIntosh, R., Wellings, C., and Park, R. (1995). Wheat Rusts: An atlast of resistance genes. CSIRO.

Mitchell, A.G., Martin, C.E. (1997). Fah1p, a Saccharomyces cerevisiae cytochrome b5 fusion protein, and its Arabidopsis thaliana homolog that lacks the cytochrome b5 domain both function in the α-hydroxylation of sphingolipid-associated very long chain fatty acids. J. Biol. Chem. 272, 28281–28288.

Moerschbacher, B.M., Vander, P., Springer, C., Noll, U., Schmittmann, G. (1994). Photosynthesis in stem rust-infected, resistant and susceptible near-isogenic wheat leaves. Can. J. Bot. 72, 990–997

Nagano, M., Takahara, K., Fujimoto, M., Tsutsumi, N., Uchimiya, H., Kawai-Yamada, M. (2012). Arabidopsis sphingolipid fatty acid 2-hydroxylases (AtFAH1 and AtFAH2) are functionally differentiated in fatty acid 2-hydroxylation and stress responses. Plant physiol. 159(3), 1138–1148.

Nekrasov, V., Wang, C., Win, J., Lanz, C., Weigel, D., Kamoun, S. (2017). Rapid generation of a transgene-free powdery mildew resistant tomato by genome deletion. Sci. Rep. 7

Olivera, P., Newcomb, M., Szabo, L. J., Rouse, M., Johnson, J., Gale, S., et al. (2015). Phenotypic and Genotypic Characterization of Race TKTTF of Puccinia graminis f. sp. tritici that Caused a Wheat Stem Rust Epidemic in Southern Ethiopia in 2013–14. Phytopathology. 105, 917–928

Omidvar, V., Dugyala, S., Li, F., Rottschaefer, S. M., Miller, M. E., Ayliffe, M., et al. (2018). Detection of race-specific resistance against Puccinia coronata f. sp. avenae in Brachypodium species. Phytopathology. 108, 1443–1454

Paradis, E., and Schliep, K. (2018). ape 5.0: an environment for modern phylogenetics and evolutionary analyses in R. Bioinformatics. 35, 526–528

Periyannan, S., Milne, R. J., Figueroa, M., Lagudah, E. S., and Dodds, P. N. (2017). An overview of genetic rust resistance: from broad to specific mechanisms. PLoS Pathog. 13

Pessina, S., Pavan, S., Catalano, D., Gallotta, A., Visser, R.G.F., Bai, Y., Malnoy, M., and Schouten, H.J. (2014). Characterization of the MLO gene family in Rosaceae and gene expression analysis in Malus domestica. BMC Genomics 15, 618

Peterson, P. D. (2001). Stem rust of wheat: from ancient enemy to modern foe. American Phytopathological Society, St. Paul, USA.

Petre, B., Joly, D. L., and Duplessis, S. (2014). Effector proteins of rust fungi. Front.Plant Sci. 5:416

Pretorius, Z. A., Singh, R. P., Wagoire, W. W., Payne, T. S. (2000). Detection of Virulence to Wheat Stem Rust Resistance Gene Sr31 in Puccinia graminis f. sp. tritici in Uganda. Plant Dis. 84, 203

Rate D.N., Greenberg J.T. (2001). The Arabidopsis aberrant growth and death2 mutant shows resistance to Pseudomonas syringae and reveals a role for NPR1 in suppressing hypersensitive cell death. Plant J. 27, 203–11

Robin, G.P., Kleemann, J., Neumann, U., Cabre, L., Dallery, J.F., Lapalu, N., et al. (2018). Subcellular localization screening of Colletotrichum higginsianum effector candidates identifies fungal proteins targeted to plant peroxisomes, Golgi bodies, and microtubules. Front. Plant Sci. 9, 562

Rushton, P.J., Somssich, I.E., Ringler, P., Shen, Q.J. (2010). WRKY transcription factors. Trends Plant Sci. 15, 247–258

Salcedo, A., Rutter, W., Wang, S., Akhunova, A., Bolus, S., Chao, S., et al. (2017). Variation in the AvrSr35 gene determines Sr35 resistance against wheat stem rust race Ug99. Science. 358, 1604–1606

Schaefer, R.J., Michno, J.M., Jeffers, J., Hoekenga, O., Dilkes, B., Baxter, I., et al. (2018). Integrating Coexpression Networks with GWAS to Prioritize Causal Genes in Maize. The Plant Cell. 30(12), 2922–2942

Singh, R. P., Hodson, D. P., Jin, Y., Lagudah, E. S., Ayliffe, M. A., Bhavani, S., et al. (2015). Emergence and Spread of New Races of Wheat Stem Rust Fungus: Continued Threat to Food Security and Prospects of Genetic Control. Phytopathology. 105, 872–884

Song J.T., Lu H., Greenberg J.T. (2004). Divergent roles in Arabidopsis thaliana development and defense of two homologous genes, aberrant growth and death2 and AGD2-LIKE DEFENSE RESPONSE PROTEIN1, encoding novel aminotransferases. Plant Cell. 16, 353–66

Staples, R., and Macko, V. (1984). Germination of Urediospores and Differentiation of Infection Structures. Pages 255–289 in: The Cereal Rusts, W. Bushnell, ed. Academic Press.

Steffenson, B. J., Case, A. J., Pretorius, Z. A., Coetzee, V., Kloppers, F. J., Zhou, H., et al. (2017). Vulnerability of Barley to African Pathotypes of Puccinia graminis f. sp. tritici and Sources of Resistance. Phytopathology. 107, 950–962

Sun, K., van Tuinen, A., van Kan, J.A.L., Wolters, A.-M.A., Jacobsen, E., Visser, R.G.F., et al. (2017). Silencing of DND1 in potato and tomato impedes conidial germination, attachment and hyphal growth of Botrytis cinerea. BMC Plant Biol. 17, 235

Sun, K., Wolters, A.-M.A., Vossen, J.H., Rouwet, M.E., Loonen, A.E.H.M., Jacobsen, E., et al. (2016). Silencing of six susceptibility genes results in potato late blight resistance. Transgenic Res. 25, 731–742

Su’udi M., Kim M.G., Park S.R., Hwang D.J., Bae S.C., Ahn I.P. (2011). Arabidopsis cell death in compatible and incompatible interactions with Alternaria brassicicola. Mol. Cells. 31, 593–601

Upadhyaya, N.M., Mago, R., Panwar, V., Hewitt, T., Luo, M., Chen, J., et al. Genomics accelerated isolation of a new stem rust avirulence gene - wheat resistance gene pair. Nat. Plants. In press.

Urnov F.D., Rebar, E.J., Holmes M.C., Zhang, H.S., Gregory, P.D. (2010). Genome editing with engineered zinc finger nucleases. Nat. Rev. Genet. 11, 636–646

Van Damme M., Andel A., Huibers R.P., Panstruga R., Weisbeek P.J., Van den Ackerveken G. (2005). Identification of Arabidopsis loci required for susceptibility to the downy mildew pathogen Hyaloperonospora parasitica. Mol. Plant Microbe In. 18, 583–92

Van Damme M., Huibers R.P., Elberse J., Van den Ackerveken G. (2008). Arabidopsis DMR6 encodes a putative 2OG-Fe(II) oxygenase that is defense-associated but required for susceptibility to downy mildew. Plant J. 54, 785–93

Van Schie, C. C., and Takken, F. L. (2014). Susceptibility genes 101: how to be a good host. Annu. Rev. Phytopathology. 52, 551–581

Vogel, J., Hill, T. (2008). High-efficiency Agrobacterium-mediated transformation of Brachypodium distachyon inbred line Bd21-3. Plant Cell Rep. 27, 471–478

Vogel, J. P., Tuna, M., Budak, H., Huo, N., Gu, Y.Q., Steinwand, M. A. (2009). Development of SSR markers and analysis of diversity in Turkish populations of Brachypodium distachyon. BMC Plant Biol. 9, 88.

Wang, X., Tang, C., Huang X., Li, F, Chen, X., Zhang G., et al. (2012). Wheat BAX inhibitor-1 contributes to wheat resistance to Puccinia striiformis. J. Exp. Bot. 63(12), 4571–4584

Wang, Y., Cheng X., Shan Q., Zhang, Y., Liu, J., Gao, C., et al. (2014). Simultaneous editing of three homoeoalleles in hexaploid bread wheat confers heritable resistance to powdery mildew. Nat. Biotechnol. 32, 947–951

Waterhouse, P., Helliwell, C. (2003). Exploring plant genomes by RNA-induced gene silencing. Nat. Rev. Genet. 4, 29–38

Wickham, H. 2016. ggplot2: Elegant Graphics for Data Analysis. Springer-Verlag, New York.

Wildermuth, M. C. (2010). Modulation of host nuclear ploidy: a common plant biotroph mechanism. Curr. Opin. Plant Biol. 13, 449–458

Yadav, I. S., Sharma, A., Kaur, S., Nahar, N., Bhardwaj, S. C., Sharma, T.R., et al. (2016). Comparative temporal transcriptome profiling of wheat near isogenic line carrying Lr57 under compatible and incompatible interactions. Front. Plant Sci. 7, 1943

Yu, G., Smith, D.K., Zhu, H., Guan, Y., and Lam, T.T.-Y. (2017). ggtree: an r package for visualization and annotation of phylogenetic trees with their covariates and other associated data. Methods Ecol. Evol. 8, 28–36

Yu, I., Fengler, K.A., Clough, S.J., Bent, A.F. (2000). Identification of Arabidopsis mutants exhibiting an altered hypersensitive response in gene-for-gene disease resistance. Mol. Plant-Microbe Interact. 13(3), 277–86.

Zambino, P.J., Kubelik, A.R., and Szabo, L.J. (2000). Gene action and linkage of avirulence genes to DNA markers in the rust fungus Puccinia graminis. Phytopathology. 90, 819–826

Zaidi, S.S., Mukhtar, M.S., Mansoor, S. (2018). Genome Editing: Targeting Susceptibility Genes for Plant Disease Resistance. Trends Biotechnol. 36(9), 898–906

Zhang, H., Yang, Y., Wang, C., Liu, M., Li, H., Fu, Y., et al. (2014). Large-scale transcriptome comparison reveals distinct gene activations in wheat responding to stripe rust and powdery mildew. BMC Genomics. 15, 898

Zhang, Y., Zhao, L., Zhao, J., Li, Y., Wang, J., Guo, R., et al. (2017). S5H/DMR6 Encodes a Salicylic Acid 5-Hydroxylase That Fine-Tunes Salicylic Acid Homeostasis. Plant Physiol. 175, 1082–1093

Zheng, Z., Mosher, S.L., Fan, B., Klessig, D.F., Chen, Z. (2007). Functional analysis of Arabidopsis WRKY25 transcription factor in plant defense against Pseudomonas syringae. BMC Plant Biol. 7, 2

Zeilmaker, T., Ludwig, N.R., Elberse, J., Seidl, M.F., Berke, L., Van Doorn, A., et al. (2015). DOWNY MILDEW RESISTANT 6 and DMR6-LIKE OXYGENASE 1 are partially redundant but distinct suppressors of immunity in Arabidopsis. Plant J. 81, 210–222

